# GRAVITY: Dynamic gene regulatory network-enhanced RNA velocity modeling for trajectory inference and biological discovery

**DOI:** 10.64898/2026.01.31.702983

**Authors:** Ziyang Miao, Zhaoyu Fang, Xingyuan Shi, Yanping Zeng, Tianrui Wu, Ruiqing Zheng, Min Li

## Abstract

RNA velocity techniques have emerged as efficient tools for unraveling the complex trajectories of cell development and differentiation. However, most of existing RNA velocity approaches are constrained by estimating transcriptional parameters for each gene in isolation and neglects the regulatory relationships among genes, which limits the ability to jointly resolve the dynamic rewiring of gene regulation and the underlying gene transcriptional kinetics across cell state transitions. To address these limitations, we present GRAVITY, a novel deep learning framework that explicitly integrates regulatory dynamics into transcriptional kinetics inference and utilizes a refined two-stage optimization strategy. Benchmarking across various simulated and real single-cell RNA sequencing datasets demonstrates that GRAVITY accurately infers both cellular and gene trajectories, along with their associated kinetic parameters. Most importantly, GRAVITY uncovers terminal cell states L5/L6 in embryonic brain development dataset. Furthermore, GRAVITY not only provides mechanistic insights by identifying the driver regulatory factors and modules governing cell fate, but also enables the systematic in silico simulation of cellular velocity changes induced by targeted regulatory perturbations.

## Introduction

Advances in single-cell RNA sequencing (scRNA-seq)^1^ have revolutionized the understanding of cellular heterogeneity by enabling high resolution transcriptome-wide profiling at the individual cell level, providing mechanistic insights into cellular development and differentiation processes. However, the requisite cell lysis step restricts temporal analysis to static snapshots, necessitating computational trajectory inference to reconstruct the continuous state transitions obscured by this technical limitation. Traditional trajectory inference methods, including PAGA^2^, Slingshot^3^, and Monocle^4, 5^, rely heavily on computing cell-to-cell similarities and typically require prior biological information such as cell type annotations, designated starting states, or manually assigned directions. Hence, researchers have begun pursuing novel strategies independent of predefined biological priors, among which RNA velocity stands as a breakthrough through systematic analysis of transcriptional dynamics^6^. By combining measurements of unspliced and spliced RNA abundances with the formulation of straightforward dynamical equations governing RNA transcription, splicing, and degradation, RNA velocity methods have not only overcome the limitations of traditional trajectory inference approaches but also attained extensive adoption in the field. Velocyto^6^ established the foundational framework for RNA velocity by leveraging a steady-state transcriptional assumption, whereby sharing the same kinetic rate parameters across all genes within individual cells. More recently, a variety of approaches have been proposed to extend the foundational framework, enabling it to capture the complex and heterogeneous nature of gene kinetics observed in real biological processes more accurately. scVelo^7^ resolves the full transcriptional-splicing kinetics through an iterative expectation-maximization (EM) algorithm, establishing a generalized dynamical system model that accurately captures transient states under non-stationary conditions. UniTVelo^8^ introduces a top-down, spliced RNA-centric framework that innovatively models transcriptional dynamics with radial basis function *RBF*-parameterized transcription rate functions, then estimates a unified latent time by iteratively aggregating gene-specific temporal information. While mathematical frameworks continuously refine the modeling of RNA velocity, recent advances in deep learning have further enhanced RNA velocity estimation by enabling data-driven modeling of nonlinear transcriptional dynamics. DeepVelo^9^ estimates the cell and gene-specific kinetic rates of transcription, splicing, and degradation using a deep neural network. CellDancer^10^ introduces a novel deep neural network-based relay velocity framework that infers cell-specific RNA velocities from neighboring cells and iteratively propagates them across whole transcriptome. VeloVI^11^ addresses RNA velocity uncertainty quantification through a probabilistic deep generative model.

Besides, several innovative approaches further transcend the classical unspliced/spliced RNA paradigm through systematic integration of additional omics and biological information. For instance, MultiVelo^12^ resolves cross-omic dynamics by incorporating chromatin accessibility with transcriptome kinetics. Dynamo^13^ integrates metabolic labeling-enabled absolute RNA velocity estimation with continuous transcriptomic vector field reconstruction. To capture specific cell cycle trajectories, DeepCycle^14^ leverages phase-specific transcriptional dynamics of core cell cycle genes, while VeloCycle^15^ unifies manifold topological learning and velocity inference using a generative model. Distinct from other methods, TFvelo^16^ replaces splicing-based RNA velocity kinetics with a TF(transcription factor)-centric framework, which explicitly incorporates the expression profiles of TFs and gene regulatory information. Although existing RNA velocity methods have demonstrated utility in reconstructing cell temporal dynamics, their effectiveness remains limited by the kinetic assumptions in transcriptional modeling, and systematic biases propagated through subsequent analytical implementations. First of all, most methods focus on optimizing each gene-level kinetic rates, whereas the characterization of cell-level velocity is achieved solely through the summation and smoothing of these individual gene rates. This bottom-up design fails to account for the dynamic changes in gene contributions to cell-level velocity across different developmental stages, branches events, and sub-lineage trajectories. On the other hand, existing methods neglect the underlying regulatory interactions between genes. Although some approaches, such as TFVelo and MultiVelo, incorporate changes in TFs’ expression and chromatin accessibility, they remain confined to single-gene analyses and do not address the dynamic rewiring of gene regulation^17^. Meanwhile, despite their predictive accuracy, deep learning-based RNA velocity methods usually suffer from the “black-box” nature of neural networks, thereby limiting the biological interpretability of inference results.

In this study, we present a deep learning framework, called GRAVITY, that overcomes those above challenges by bridging transcriptional dynamics with regulatory mechanism through joint modeling of RNA velocity and gene regulations. By processing the abundance of unspliced and spliced RNA through a regulatory-aware transformer architecture, GRAVITY decodes context-specific gene regulatory dynamics driving cell state transitions at single-cell resolution. Unlike previous approaches emphasizing estimating RNA velocity for individual gene, GRAVITY implements a two-stage optimization strategy: cell-wise and gene-wise optimization. In the cell-wise phase, GRAVITY optimizes cell-level RNA velocity vectors informed by its neighbor cells motivated by previous relay velocity model^10^. Subsequently, in the gene-wise stage, these refined cellular directions serve to constrain the inference of individual gene velocity kinetics. This two-stage strategy resolves the inherent compromise in multi-gene velocity models between individual gene transcriptional dynamics and cell-level velocity vector coherence. Benchmark analyses across various simulated and real cell lineage datasets validates GRAVITY’s capabilities and generalization of step-by-step resolving cell-level state transitions and gene-specific velocity inference. We apply GRAVITY to well-characterized biological systems: pancreatic endocrinogenesis and embryonic brain development (E18) in mice. Across these datasets, we demonstrate GRAVITY’s ability to accurately recapitulate the context-specific regulatory factors and dynamic regulatory relationships that govern cell development and fate determination. In embryonic brain development dataset, GRAVITY is the unique method that accurately identifies two terminal developmental states—the Deeper Layer and the Upper Layer. By analyzing gene contributions, we also identify and validate the two substates, L5 and L6 in the Deeper Layer. Furthermore, we develop an in silico gene perturbation strategy that implements systematic in silico perturbation of lineage-controlling TFs across pancreatic endocrinogenesis lineages. Experimental results also demonstrate that GRAVITY accurately simulates the expected shifts in cell fate trajectories after gene perturbations. All these results show that GRAVITY provide more accurate estimation of cell-specific transcriptional kinetics and decoding the underlying regulatory mechanisms governing cell-fate determination via context-aware GRN inference.

## Results

### Overview of GRAVITY

Motivated by evidence that regulatory interactions between genes and their dynamic rewiring are critical to cellular development and differentiation^17, 18^ (Fig. 1a), we designed GRAVITY, an interpretable method for RNA velocity inference that explicitly integrates gene regulatory relationships. The overall model architecture of GRAVITY is shown in Fig. 1b. GRAVITY firstly takes the unspliced, spliced abundance matrices and 2D manifold of cells (UMAP) as input. Then, the model includes a regulatory network aware module that masks cross-attention with the NicheNet^19^ prior regulatory network, thereby elucidating regulatory relationships between genes (Fig. 1c), derives unified cell representations via a kinetic parameter inference module and optimizes the model using a dedicated loss module.

**Figure 1.**
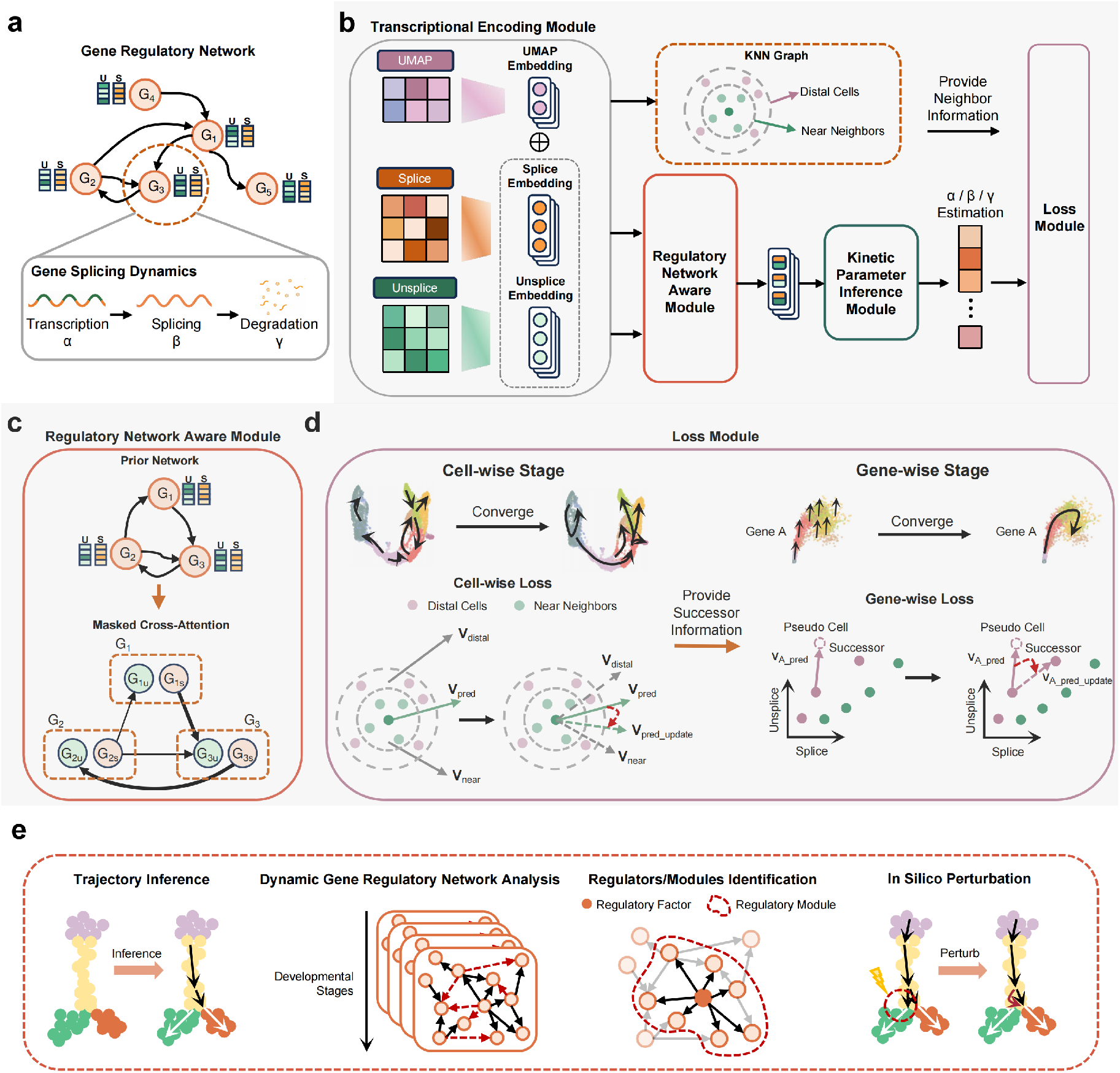
An overview of GRAVITY. **a**, Modeling of transcriptional kinetics. Within a specific cell, the regulatory relationships among genes influence individual gene’s transcription (α), splicing (β), and degradation (γ) rates. **b**, GRAVITY takes unspliced/spliced abundance of genes and cell embeddings (UMAP) as input, captures regulatory relationships via regulatory network aware module, and estimates cell- and gene-specific kinetic parameters through a shared MLP. **c**, Regulatory network aware module resolves the dynamic GRN by leveraging cross-attention matrices between the spliced representations of regulatory genes and the unspliced representations of target genes, which are masked by a background GRN. **d**, The two-stage optimization strategy of GRAVITY first optimizes the overall cellular velocity and trajectory, then refines gene-specific kinetic rates based on optimized cellular trajectory. **e**, GRAVITY facilitates a comprehensive suite of downstream analyses, including trajectory inference, dynamic network analysis, identification of regulators and modules, and in silico perturbation experiments, thereby providing substantial biological insights.

Unlike most of previous single-gene velocity inference approaches, GRAVITY is a whole-transcriptome model that prioritizes the reconstruction of global cellular trajectories and system-wide velocity inference. While this design provides biological insights of cellular dynamics, it presents inherent trade-offs in optimizing gene-specific kinetic parameters. To address this issue, GRAVITY employs a two-stage optimization strategy in loss module (Fig. 1d). In the cell-wise stage, the model infers RNA velocity for each cell jointly across all genes and ensures the consistency between the predicted velocity vector and their local neighborhood directions. During the gene-wise stage, GRAVITY leverages the cell-derived global velocity vectors as reference manifolds and fixes parameter updates exclusively to the last three layers of the shared MLP to optimize gene-specific RNA velocity inference. In summary, GRAVITY not only achieves superior overall performance in RNA velocity inference, but also enables cell-fate-centric downstream analysis—including dynamic GRNs reconstruction and mining, and in silico perturbation studies—through the interpretable attention mechanism (Fig. 1e).

### GRAVITY delivers superior trajectory inference and robustness to prior network noise on simulated datasets

To evaluate the performance of GRAVITY, we first used four simulated datasets generated by SERGIO^20^ and conducted benchmark experiments against three typical RNA velocity methods, including the dynamic-modeling framework scVelo^7^, the deep learning-based framework CellDancer^10^, and the regulatory-based framework TFvelo^16^. SERGIO is capable of utilizing stochastic differential equations (SDEs) to model the dynamics of both unspliced and spliced gene transcripts as functions of the fluctuating levels of their regulatory transcription factors (TFs), as determined by a predefined GRN. The four simulated datasets exhibited a trajectory topology of increasing complexity, including linear, bifurcating, trifurcating, and quadrifurcating (Fig. 2a). Each dataset contains 100 genes, including 10 TFs, and the underlying gene regulatory network, but differs in the number of cells. The RNA velocity inference results obtained by different methods for all datasets are shown in Fig. 2. Representatively, in all four datasets, GRAVITY demonstrated superior trajectory inference performance and exhibited higher consistency with the ground truth (Fig. 2b-e). In comparison, scVelo exhibited mild backflow at the end of developmental trajectories, while both CellDancer and TFvelo showed inconsistent initiation of the trajectory. Specifically, TFvelo relies on the expression level of TFs rather than genes’ unspliced information. An overlap analysis between TFvelo-selected transcription factors and the ground-truth set showed limited concordance in the trifurcating setting and moderate concordance in the bifurcating setting (Fig. S2), suggesting that its performance is sensitive to the transcription-factor selection strategy. However, GRAVITY provided a clearer inference of the three differentiation branches in the trifurcating dataset compared to scVelo, CellDancer, and TFvelo (Fig. 2d). To provide a quantitative assessment of these methods, we proposed a Branching-aware Trajectory Consistency metric for RNA velocity, termed BATC (Fig. S1). This new metric mitigates the bias caused by cell states’ boundary mixture and provides more accurate evaluation of the consistency between cells’ velocity vector and the real cross-stage trajectory (Fig. S3). As expected, GRAVITY achieved a higher BATC score than scVelo, CellDancer, and TFvelo across the four datasets. On average, GRAVITY outperformed the next-highest scoring method, scVelo, by 5.3% (Fig. 2f). To quantify the contribution of the prior network, we ablated it and evaluated GRAVITY_naive. Despite this removal, GRAVITY_naive led three of four simulated datasets, with the full GRAVITY retaining the highest overall scores (Fig. 2f). Nevertheless, incorporating the prior GRN improved velocity prediction accuracy in most cases, indicating that reliable prior gene regulatory relationships can reduce confounding indirect regulations and thereby enable more accurate inference of RNA velocity. Considering the difficulty in obtaining accurate prior GRNs in real-world applications, we further assessed the impact of network noise on RNA velocity inference. We incrementally added noise edges to the prior GRN (from 1-to 5-fold of original edges), performed 10 random replicates for each setting, and evaluated the BATC of GRAVITY across four simulated datasets. GRAVITY exhibits considerable robustness to the noise in the prior GRN. To quantify this, its performance was evaluated across four datasets under various simulated noise levels (Fig. 2g). GRAVITY maintained stable performance, achieving a mean BATC of 0.267 ± 0.122 on Dataset A and 0.624 ± 0.058 on Dataset B. On the more challenging datasets C and D, the method achieved BATC scores of 0.597 ± 0.038 and 0.518 ± 0.044, respectively. The low standard deviation across most scenarios underscores the model’s stability against perturbations in the input network.

**Figure 2.**
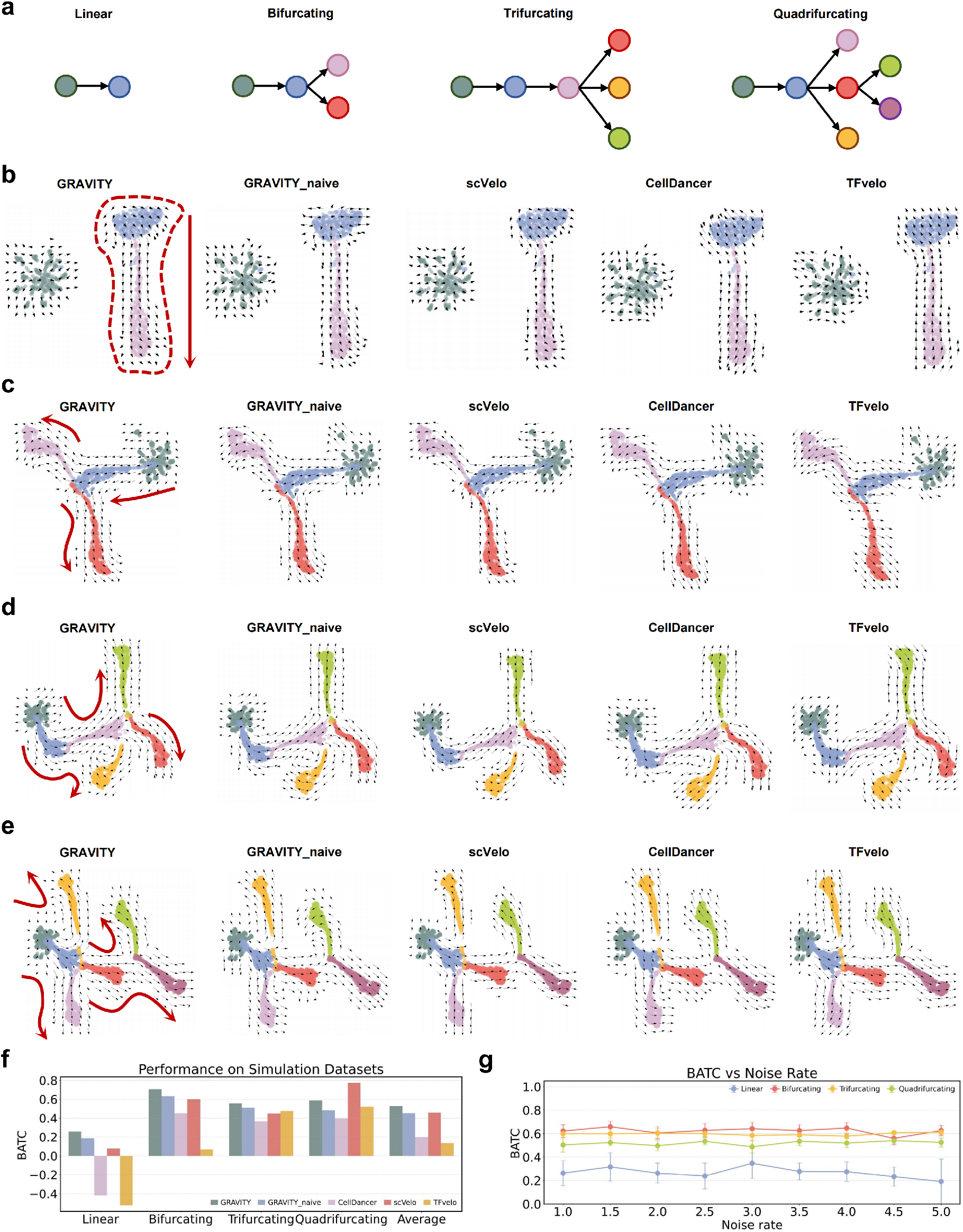
Benchmarking GRAVITY on simulated datasets. **a**, Schematic diagrams of four simulated trajectory backbones with increasing complexity (linear, bifurcating, trifurcating and quadrifurcating) in thesimulated datasets. **b–e**, The comparison among inferred celluar dynamics of GRAVITY, GRAVITY_naive, scVelo, CellDancer and TFvelo across the four simulated datasets. The red arrows in the GRAVITY panels indicate the ground-truth trajectory direction. **f**, Evaluation of the five methods using the BATC metric. **g**, Robustness of GRAVITY’s performance in response to increasing noise levels in the background GRN.

### GRAVITY accurately infers cellular dynamics, pseudotime and RNA kinetic parameters on real-world datasets

For a comprehensive evaluation of GRAVITY’s performance in real-world scenarios, we applied it and other competitive methods to two previously reported datasets, including human forebrain development dataset^6^ and the scEU-seq cell-cycle dataset of RPE-1 cells^21^. The human forebrain developmental dataset defines a clear linear trajectory across seven continuous cellular states, transitioning from radial glia to differentiated mature glutamatergic neurons^6^. At the cellular level, GRAVITY accurately infers the velocity pattern along this continuous trajectory. In contrast, CellDancer exhibits velocity reversals at both termini, and scVelo also demonstrates erroneous directional inference throughout the intermediate trajectory (Fig. 3a). Besides the global analysis of cellular progression directionality, we compared the inferred velocities of individual genes using gene-specific phase portraits To illustrate the complexities inherent in multi-kinetic gene behavior, we selected *CNTNAP*2, which is essential for neuronal development and synaptic function^22^, and *GNAO*1, a pivotal regulator of G protein signaling^23^. Based on the expression of unspliced and spliced mRNA, both genes underwent relatively complete induction and repression phases. Only the velocities inferred by GRAVITY accurately aligned with the cellular developmental trajectory. Notably, scVelo unexpectedly omitted *GNAO*1 from its cellular velocity calculations, resulting in no inferred velocity vector for this gene, while CellDancer’s inferred velocity direction did not match the actual developmental trajectory. Since TFvelo employs TF-centric phase portraits, direct comparison is infeasible. However, we clearly observed that the inferred velocities for *CNTNAP*2 and *GNAO*1 were ambiguous between the different developmental states.

**Figure 3.**
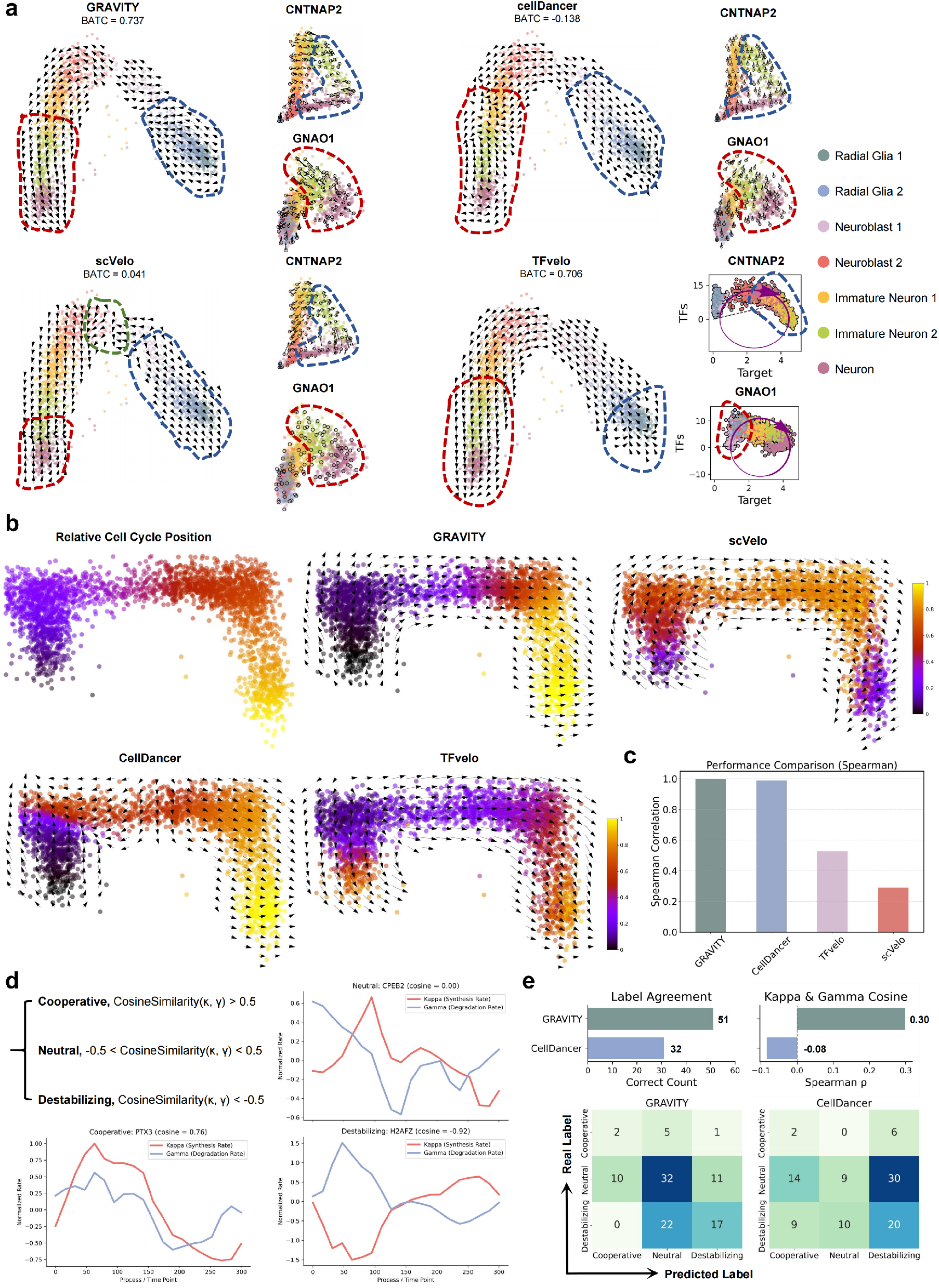
Benchmarking GRAVITY on real scRNA-seq datasets. **a**, Cellular and two representative gene dynamics (*CNTNAP*2, *GNAO*1) inferred by GRAVITY, scVelo, CellDancer, and TFvelo on the human forebrain development dataset. **b**, Comparison of pseudotime and cellular dynamics inferred by the four methods across the scEU-seq cell-cycle dataset. **c**, Spearman’s rank correlation (ρ) between inferred pseudotime and true cell-cycle stages. **d**, Three distinct three turnover strategies of mRNA regulation during the cell cycle based on measured κ and γ the scEU-seq–provided κ and γ. **e**, Evaluation of label concordance between ground-truth mRNA regulation patterns and those predicted by GRAVITY and CellDancer, along with correlations between inferred and true kinetic parameters (κ, γ).

Next, we investigated the predicted cell velocities and gene kinetic parameters using a metabolic labeling (scEU-seq^21^) cell-cycle dataset from RPE-1 cells. We began by visualizing the cell velocities and corresponding pseudotime, as derived by GRAVITY and other methods (Fig. 3b). These velocities and pesudotime were mapped onto the relative position along the cell cycle, which was determined using the Geminin-GFP and Ct1-RFP signals from the FUCCI system. The results indicate that GRAVITY achieves the strongest consistency with the actual cell cycle processes in both trajectory and pseudotime inference; scVelo likewise produces accurate trajectories, while CellDancer and TFvelo exhibited notable deficiencies in the initial and intermediate stages, respectively. Moreover, GRAVITY achieved the strongest Spearman correlation between its inferred pseudotime and the 290 provided cell cycle phases (GRAVITY:0.999, CellDancer:0.988, TFvelo:0.527, scVelo:0.290, Fig. 3c). Another major advantage of the scEU-seq technique is its ability to measure the synthesis and degradation of labeled RNA within a defined period and experimental manner. This capability provides a direct and reliable benchmark for evaluating the accuracy of inferred gene kinetics parameters. Original study^21^ defined three turnover strategies of mRNA regulation during the cell cycle based on the cosine similarity between measured synthesis and degradation rates: Cooperative (cosine > 0.5), Neutral (-0.5 < cosine < 0.5), and Destabilizing (cosine < -0.5) (Fig. 3d). Biologically, evaluating the accuracy of inferred mRNA regulatory patterns provides greater insight than directly comparing the predictive accuracy of gene kinetic parameter values. Using the same definition, we classified the top 100 genes most correlated with cell cycle phases based on the transcription rate κ and degradation rate γ estimated by computational methods (Fig. 3e). For methodological reasons, scVelo and TFvelo were excluded from this evaluation, as scVelo does not infer gene kinetic parameters at the single-cell level and TFvelo does not estimate transcription and degradation rates. In comparison with CellDancer, GRAVITY correctly classified 51 of the 100 genes, outperforming CellDancer’s 32, with the greatest improvement observed in the Neutral category. We further calculated the Spearman correlation between the scEU-seq-derived cosine similarity and the computationally inferred cosine similarity for all genes (Fig. 3e). GRAVITY achieved higher correlation than CellDancer (0.30 vs. -0.08), highlighting GRAVITY’s superior ability to capture gene kinetic trends and mRNA regulatory patterns.

### GRAVITY reveals pancreatic cell fate determination, regulatory factors and in silico perturbed transdifferentiation predictions

We applied GRAVITY to the endocrine development of the mouse pancreas profiled from embryonic day 15.5 (E15.5)^24^, to demonstrate its capability of inferring transient lineages and resolving the dynamics of regulatory network rewiring. This dataset comprises four major endocrine cell fates—glucagon-producing alpha cells, insulin-producing beta cells, somatostatin-producing delta cells, and ghrelin-producing epsilon cells—as well as a cycling subpopulation of ductal cells and endocrine progenitor cells^25^. For cellular velocity inference, GRAVITY accurately resolved both velocity vectors for the four endocrine differentiation processes and the directionality of the cycling subpopulations (Fig. 4a). Moreover, comparison of cell velocities before and after the second gene-wise stage revealed that the second stage training further corrected the absolute magnitude of cell velocities (Fig. S4a). Notably, magnitudes in all four mature endocrine cell types decreased significantly, falling below that of the transient pre-endocrine phase. This observation aligns with reduced nascent RNA synthesis at terminal developmental stages, as well as the equilibration of unspliced and spliced RNA ratios, resulting in near-zero RNA velocity magnitudes. Having established cellular velocity, we also examined whether GRAVITY could resolve gene-specific velocity patterns for phase portrait fitting. We selected three representative genes (*Bicc*1, *Rb f ox*3, *R f x*6) for comparison and visualized the gene velocity vectors before/after the second gene-wise stage (Fig. 4b). For *Bicc*1, previous study have validated that it is specifically expressed in pancreatic progenitor and ductal cells^26^, but is absent from mature endocrine cells. While both GRAVITY and scVelo detected the repression phase-specific kinetic pattern, GRAVITY-derived velocity more accurately recapitulated the overall phase portrait dynamics. Furthermore, the pancreatic endocrine development related gene *Rb f ox*3 and *R f x*6 exhibit biphasic expression dynamics during differentiation (Fig. S4b). Only GRAVITY accurately resolved both the induction and repression phases of these genes, scVelo primarily detected induction phases, and CellDancer was limited to identifying repression phases. Notably, comparative analysis of gene velocity vectors before/after the second training stage demonstrates that GRAVITY effectively optimizes individual gene kinetic rates by leveraging refined cellular velocity constraints.

**Figure 4.**
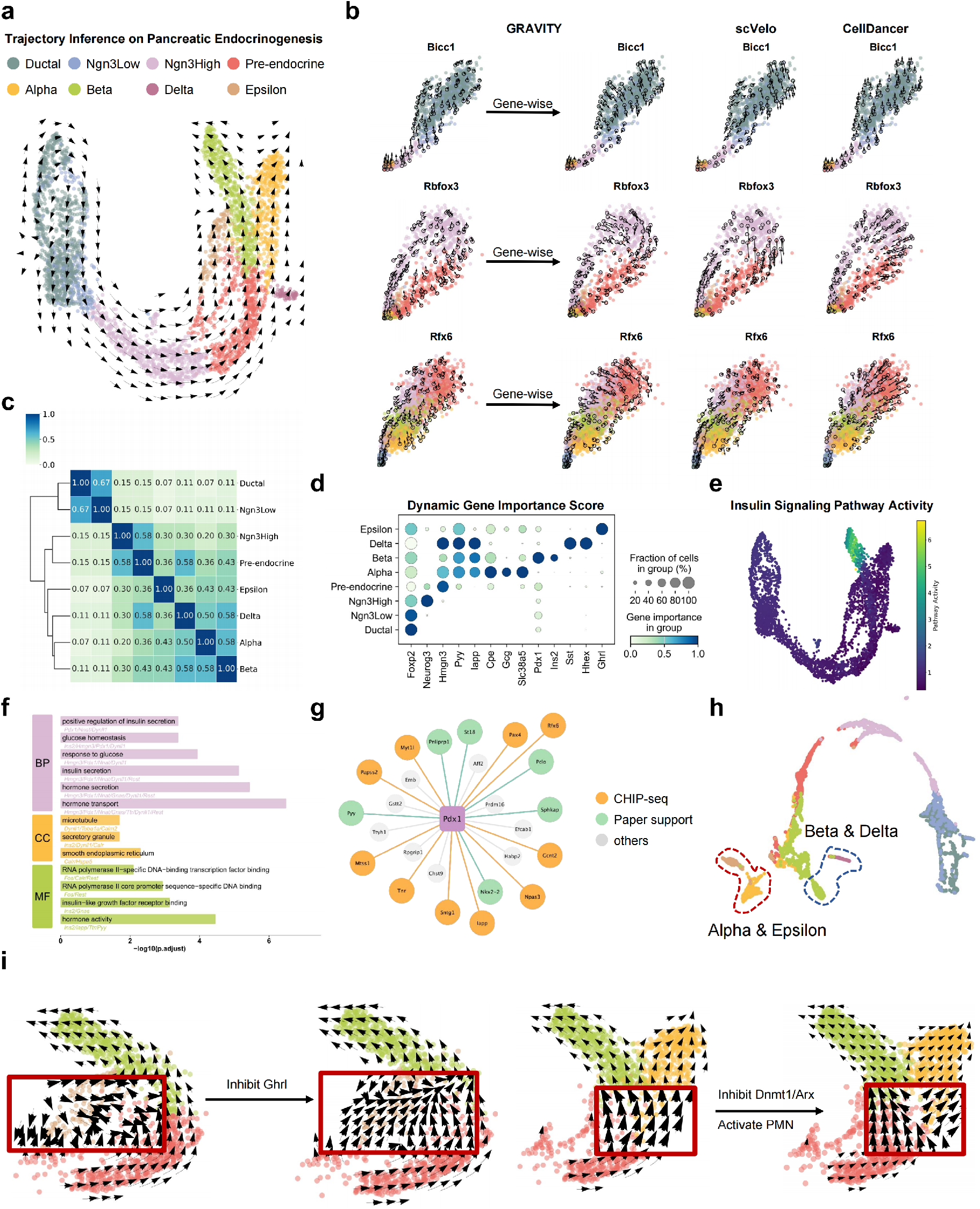
Uncovering dynamic regulatory patterns during mouse pancreatic development. **a**, Cellular dynamics inferred by GRAVITY. **b**, The phaseportrait fittings for three representative genes (*Bicc*1, *Rb f ox*3, and *R f x*6) before and after genewise training against scVelo and CellDancer. **c**, Reconstruction of pancreatic cell lineages based on inferred GRN similarity. **d**, Dot plot displaying the importance scores of well-characterized regulatory marker genes. **e**, Pathway activity scores for the insulin signalling pathway. **f**, GO enrichment analysis of cell-specific genes identified by GRAVITY, demonstrating a significant enrichment for key Beta cell biological function. **g**, Regulatory module for *Pdx*1. Orange nodes are target genes validated by CHIP-seq databases; green nodes are genes reported in related literature associated with *Pdx*1 or beta cell functions. **h**, GRAVITY embedding delineates a lineage trajectory highly consistent with the known endocrine cell developmental hierarchy. **i**, In silico perturbation shows two lineage shifts: *Ghrl* suppression redirects ε cell differentiation toward beta cells (left), and pseudo-reprogramming shifts α cell trajectories toward β cells (right).

To demonstrate GRAVITY’s capacity to resolve underlying changes in gene regulatory patterns, we then extracted per-cell attention matrix as weighted regulatory network to investigate biological insights. First, we constructed cell-type-specific regulatory networks, termed GRN_1‰_, by averaging attention weights across different cell types and retaining the top 1‰ of regulatory edges. For regulatory gene sets comprising each GRN_1‰_, we applied pairwise Jaccard similarity coefficients, followed by hierarchical clustering, to quantify regulatory similarity across cell types. Notably, this hierarchical regulatory similarity accurately captured the developmental lineage of pancreatic endocrine cells (Fig. 4c). Then, we defined gene importance scores and pathway importance scores based on the strength of regulatory relationships to identify lineage-dependent regulatory activity. GRAVITY identified critical regulation factors with clear developmental-lineage-specific regulatory activities (Fig. 4d), including major lineage-defining TFs in early development stage (Ductal/Ngn3Low cells, *Foxp*2)^27^, Ngn3High cells (*Neurog*3)^28^, endocrine maturation (*Hmgn*3, *Pyy* and *Iapp*)^29–31^, alpha cells (*Cpe, Gcg* and *Slc*38*a*5)^32, 33^, beta cells (*Pdx*1 and *Ins*2)^34, 35^, delta cells (*Sst* and *Hhex*)^36^ and epsilon cells (*Ghrl*)^37^. Similarly, the importance of insulin signaling pathway (KEGG:04922), as an beta-cell–specific functional pathway, exhibits a transition from an inactive/repressed state to significant activation (Adjusted p-value<10^−100^ Fig. 4e). Taking beta cells as an example, we prioritized the top 15 regulatory genes from GRN_1‰_ by the gene importance score and conducted GO enrichment analysis (Table. S1). GO enrichment result revealed highly significant associations (Adjusted p-value< 0.05) with fundamental beta-cell functions, including regulation of insulin secretion and glucose homeostasis (Fig. 4f). Same observations were found in Alpha cells (Fig. S5). For each TF, GRAVITY could also uncover its cell-type-specific target genes and regulatory program. To validate regulatory interaction prediction, we focused on the master regulatory factor *Pdx*1 in beta cells (Fig. 4g). Among its top 25 predicted targets by GRAVITY, 60% (15/25) have already been validated by ChIP–seq experiments from the Cistrome database^38^ and previous literature(Table. S2). In addition to gene regulation captured by the attention mechanism, the cellular embeddings derived by GRAVITY demonstrate higher biological consistency with experimentally defined developmental hierarchies (Fig. 4h). Specifically, these embeddings accurately recapitulate the *Pax*4/*Arx*-mediated cross-inhibitory interaction governing endcrine progenitor specialization^39, 40^, outperforming original visualization (Fig. 4a) in resolving the bifurcation into alpha/epsilon versus beta/delta sublineages.

Building upon the investigation into regulatory mechanisms modelling, we further proposed an in silico strategy to simulate developmental trajectory alterations upon TF perturbation by down-or up-scaling the expression of the targeted gene(s) in both upslice/splice transcriptomes. In parallel, we remove the perturbed gene(s) and all incident edges from the prior network, then re-evaluate the model to quantify trajectory shifts (Methods). The GRAVITY perturbation simulation results were then compared with findings from previous biological studies. First, we demonstrated GRAVITY’s in silico perturbation analysis with the *Ghrl* knockout (KO) simulation, targeting the master regulatory factor of epsilon cells. It was observed that the terminally differentiated state of epsilon cells was lost after *Ghrl* KO, instead displaying a notable tendency of further differentiation into beta cells (Fig. 4i). This finding aligns with observations in ghrelin gene knockout mice, where the absence of ghrelin signaling notably shifts the balance of endocrine progenitor cell differentiation towards other cell types, particularly increasing beta cell mass^41^. We next evaluated five additional regulatory genes that have previously been reported as key factors in the transdifferentiation between alpha and beta cells: *Dnmt*1, *Arx*, and the PMN genes (*Pdx*1, *Ma f a*, and *Neurog*3)^42^. Since the loss of *Arx* and *Dnmt*1 and the overexpression of PMN genes are known to promote alpha-to-beta cell transdifferentiation, we conduct to inhibit *Dnmt*1 and *Arx* and increase PMN gene expression. This phenomenon was partially validated by in silico perturbation experiments: in naïve alpha and beta cells, a subset of cells that initially differentiated towards the alpha cell lineage shifted towards the beta cell lineage (Fig. 4i). Overall, these results not only demonstrate GRAVITY’s effectiveness in recovering developmental regulatory mechanisms, but also confirm prior knowledge of cell fate regulation during pancreatic development. Furthermore, systematic simulations highlight GRAVITY’s capability for objective and reliable in-silico gene perturbation analysis.

### GRAVITY discovers embryonic brain development trajectories and captures potential regulatory patterns

To further evaluate GRAVITY’s capability in resolving cell fate commitment, we reanalyzed a 10x Multiome dataset of neocortical neurogenesis from the embryonic E18 mouse brain^12^. This dataset delineates that radial glial (RG) cells undergo asymmetric divisions to generate neurons and intermediate progenitor cells (IPCs). Subsequently, IPCs proliferate within the subventricular zone (SVZ), generating secondary neuronal cohorts. This process underlies the diversification of developing projection neurons into distinct subtypes, forming the basis for the neocortex’s organization into functionally specialized layers and areas. Specifically, early progenitor pools give rise to subplate (SP) and deeper-layer (L5–6) neurons, whereas later progenitor pools generate upper-layer (L2–4) neurons^43^. Motivated by this developmental scheme, we compared GRAVITY with other RNA-velocity methods that rely solely on scRNA-seq data. In these comparisons, GRAVITY’s cell-velocity and pseudotime inferences indicated that deeper-layer and upper-layer neurons represent distinct developmental branches rather than a precursor–successor relationship within a single branch (Fig. 5a). In contrast, scVelo and TFvelo failed to identify the Upper Layer as the other developmental endpoint. Although CellDancer identified the Upper Layer endpoint, its pseudotime estimation was inconsistent and exhibited unexpected backflows at the end of the Deeper Layer lineage. Even Multivelo, which incorporating additional scATAC-seq data, was also unable to identify the Upper Layer endpoint (Fig. 5b). Moreover, the visualization of GRAVITY-derived cell embeddings clearly revealed the bifurcation of these two branches within the V-SVZ group, thereby confirming the consistency between inferred velocities and cell biological similarities (Fig. S6b). Another notable observation was that within the deeper neuronal layer, the cell velocities inferred by GRAVITY also converged towards two distinct sub-terminal states. Based on the expression of marker genes *Bcl*11*b* in L5 subcerebral projection neurons and *Foxp*2 in L6 corticothalamic projection neurons, these sub-terminal states were assigned to L5 and L6 neurons^44^, respectively (Fig. 5c). In summary, the above findings indicate GRAVITY delineates cell fate commitment with greater accuracy and provides additional fine-grained insights.

**Figure 5.**
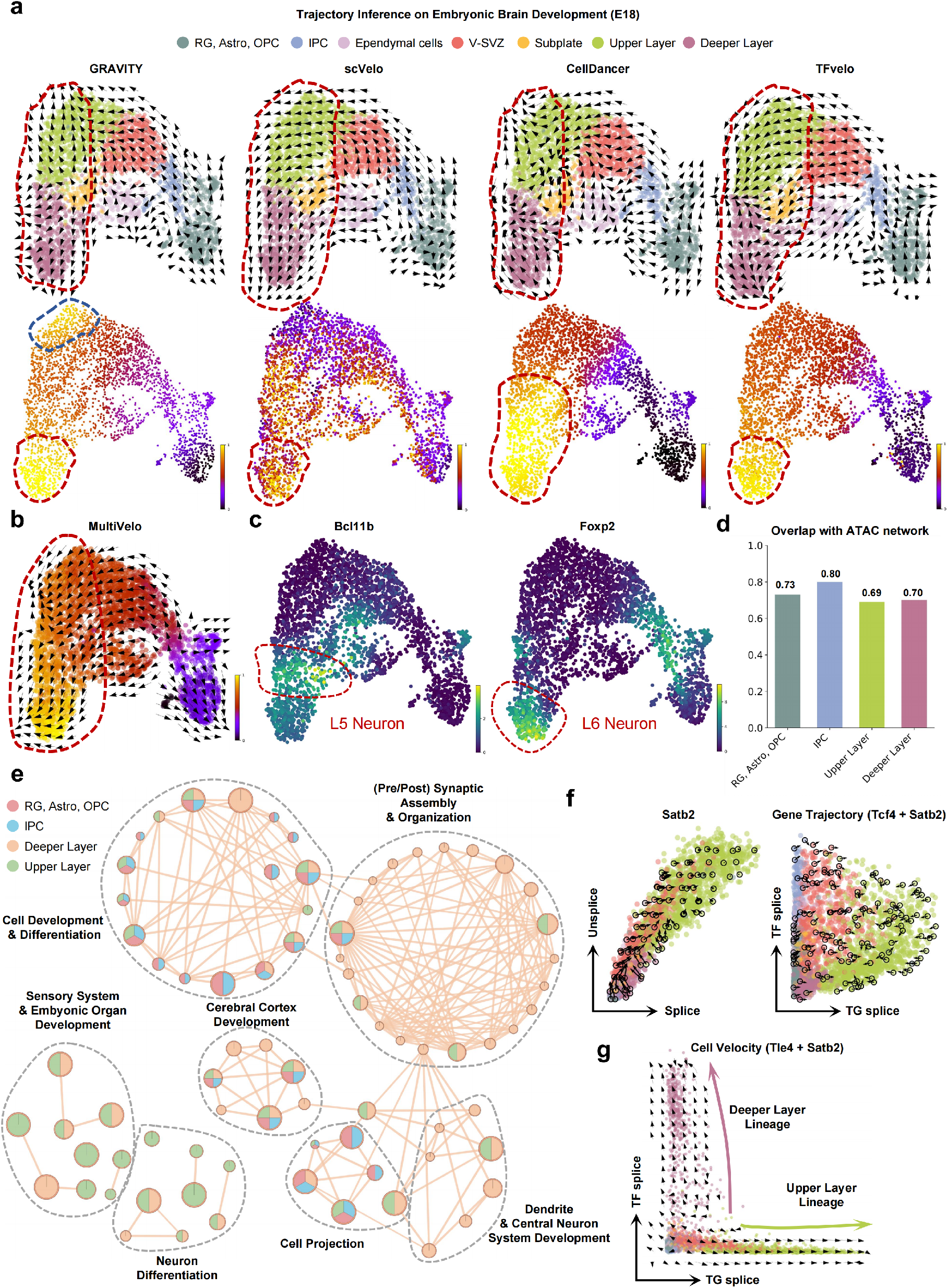
Identifying embryonic brain developmental lineages and terminal states. **a**, The cellular dynamics and corresponding pseudotime inferred by GRAVITY, scVelo, CellDancer, and TFvelo on the embryonic brain dataset. **b**, Trajectory inference by MultiVelo integrating scATAC-seq and scRNA-seq data. **c**, Gene expressions of *Bcl*11*b* and *Foxp*2, revealing potential L5 and L6 subpopulations within the Deeper Layer cells (red boxes). **d**, Accuracy of top 100 cell-type-specific regulatory edges identified by GRAVITY, validated against relationships inferred from scATAC-seq using ChromVAR. **e**, Multivariate GO enrichment analysis of identified regulatory modules across four cell types. Nodes represent BP terms, node size corresponds to gene set size, and pie colors indicate related cell states. **f**, Left panel: gene phase portrait of *Satb*2. Right panel: TF/TG-aware gene phase portrait of *Tc f* 4 and *Satb*2. **g**, Cell velocity delineated by two layer-specific transcription factors, *Tle*4 (Upper Layer) and *Satb*2 (Deeper Layer), respectively.

Likewise, we also evaluated the capability of GRAVITY to resolve the dynamics of regulatory rewiring during neocortical neurogenesis. Benefiting from the scATAC-seq data provided by the 10x Multiome technology, we first validated the accuracy of GRAVITY in inferring cell type–specific GRNs. For each cell type, the accessible chromatin peaks were identified, and peak-induced TF–TG regulatory relationships were established as the gold standard using ChromVAR^45^ and JASPAR TF motif information^46^. Cell type-specific regulatory networks inferred by GRAVITY were derived by averaging cross-attention matrices within each cell type. The top 100 highest-weight regulatory edges were then selected for validation (Fig. 5d). The accuracy for four cell types (early cell types: Radial Glia and IPC; mature cell types: Upper Layer and Deeper Layer neurons) was higher than 0.69, with the highest accuracy of 0.80 observed in IPC cells. To identify the heterogeneity of biological processes, we identified the top 30 key regulatory genes as well as top 3 TGs for each cell type based on gene importance ranking, then conducted GO enrichment analysis on the gene sets for each cell type. The multivariate enrichment results map was generated by the ActivePathways tool^47^ (Fig. 5e, Fig. S6a). Enrichment analysis revealed that early cell types were involved in cell developmental and differentiation, such as glial cell differentiation, regulation of gliogenesis, regulation of neuron projection development, etc. In contrast, terminal Upper Layer and Deeper Layer neurons exhibited enrichment for more specific functions, including neuron differentiation, regulation of synapse assembly, embryonic morphogenesis, and so on. In particular, GRAVITY also captured the distinct biological programs between these two layers, revealing that Upper Layer neurons are involved in processes related to sensory system development and neuron migration, whereas Deeper Layer neurons exhibit transcriptional programs associated with neuron axongoenesis and excitatory synapse assembly.

Further, we investigate whether the intergation of regulatory network facilitates the inference of gene kinetic. We proposed a TF/TG-aware gene phase portrait and selected *Satb*2, a gene primarily expressed in upper cortical layers with its key regulator *Tc f* 4 (also identified by GRAVITY) as an example (Fig. 5f). In the TF/TG phase portrait, *Satb*2’s velocity displayed a more clockwise trajectory relative to its original unspliced/spliced representation.

Pseudotime expression trends also clearly demonstrated a regulatory phase delay between *Tc f* 4 and *Satb*2 (Fig. S6c). These findings indicate that incorporating reliable regulatory factors enhance the inference of whole-phase gene velocities by providing richer information and simultaneously alleviating the influence of unspliced mRNA sparsity and noise. Finally, we selected *Tle*4 and *Satb*2, two TFs specific to upper and Deeper Layer neurons respectively, based on their gene importance scores. When cell velocity vectors inferred merely from these two genes (Fig. 5g), were mapped onto their spliced expression axes, we found that these two critical regulators could clearly distinguish two developmental branches of upper and Deeper Layer neurons. This result also indicates that GRAVITY effectively captured intrinsic regulatory patterns associated with developmental processes.

## Discussion

In this study, we present GRAVITY, a novel deep learning framework that integrates gene regulatory mechanisms for a robust inference of RNA velocity. In contrast to previous methods that focus on individual gene inference, GRAVITY achieves a transcription- and system-level estimation of RNA velocity. This system-level advancement is realized through three key innovations: First, in the context of gene kinetic modeling, GRAVITY generalizes the established individual-gene kinetic model to the system level by explicitly incorporating gene regulatory relationships. This is achieved by a regulatory-aware cross-attention module to capture the dynamic regulatory relationships informed by a prior GRN. Second, in RNA velocity inference, GRAVITY utilizes a top-down, two-stage optimization strategy. The initial stage optimizes for the global cellular RNA velocity, and this global velocity then serves as a reference for the next stage, which refines the kinetic parameters for individual genes. Third, owing to its inherent biological interpretability, GRAVITY not only yields mechanistic insight into the critical regulatory factors and modules governing cell fates, but also enables the systematic simulation of changes in cellular velocity induced by targeted perturbations of critical regulatory factors.

Our comprehensive benchmark tests on both simulated and real-world datasets demonstrated that GRAVITY has superior performance against existing RNA velocity inference methods, particularly in the accurate estimation of global cellular velocity, gene-specific velocity, and kinetic parameters. Notably, GRAVITY uncovered previously uncharacterized developmental branches and terminal cell states within Embryonic Brain Development and TAN polarization, highlighting its ability to refine complex development and differentiation trajectories. Furthermore, by analyzing the dynamic GRNs reconstructed by GRAVITY, we delineated the regulatory factors and modules that orchestrate cell fate decisions, and identified the rewiring of regulatory relationships across diverse biological processes, including Pancreatic Development, Embryonic Brain Development, and TAN polarization. Building upon these findings, we further validate the effects of these regulatory factors through systematic perturbation simulation in silico, without additional biological experiments. This application extends RNA velocity from trajectory reconstruction to prospective prediction.

Despite the demonstrated advantages of GRAVITY, there remain some limitations and opportunities for improvement. First, the perturbation simulation is strictly confined to the observed expression space, which means that cell states not present in the original scRNA-seq data cannot be analyzed. Second, the Regulatory Network Inference module relies on calculating gene-to-gene cross-attention. This process scales significantly with the number of genes, presenting a computational bottleneck that demands substantial GPU memory and results in reduced computational speed. Furthermore, GRAVITY currently only accounts for intrinsic gene regulatory relationships when inferring RNA velocity, neglecting the influence of other crucial biological processes such as chromatin accessibility, DNA methylation, or extracellular cell communication. Integrating single-cell multi-omics data or spatial transcriptomic information also presents a promising avenue for future enhancement.

## Methods

### Collections and preprocessing of scRNA-seq datasets

All scRNA-seq datasets, including simulated and real, were derived from publicly available repositories or studies. Four simulated datasets obtained from the SERGIO GitHub repository, each dataset contains 100 genes and 10 TFs^20^. For fairness, GRAVITY and all other methods used the same settings, without any additional preprocessing steps applied to the real datasets. The human forebrain dataset consists of 1,720 cells, with highly variable genes (HVGs) automatically selected by calculating expression dispersion. To better evaluate the interpretability of gene regulation, the pancreatic dataset retains 2,000 HVGs. The 10x embryonic brain dataset, consisting of 3,365 cells, was preprocessed identically to the TFvelo.

### Modeling transcriptional dynamics with gene regulation

Let *u*_*i j*_(*t*) and *s*_*i j*_(*t*) denote the unspliced and spliced abundances of gene *i* in cell *j* at a specific timepoint *t*, the classical kinetics of transcriptional dynamics could be summarized by the following ordinary differential equations:

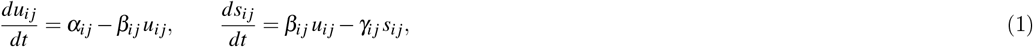

Here, the kinetic parameters α_*i j*_, β_*i j*_, and γ_*i j*_ denote the transcription, splicing, and degradation rates of gene *i* in cell *j*. These kinetic parameters can be estimated using a closed-form or analytic solution (e.g., likelihood/expectation-maximization methods, deep neural network)(Supplementary Notes). To better characterize the intrinsic transcriptional complexity, GRAVITY employs a novel deep neural network (DNN) architecture to predict these kinetic parameters. A major limitation of this transcriptional dynamics model is the assumption that the individual gene kinetic parameters are determined solely by its own unspliced and spliced abundances, which neglects crucial regulatory and interaction relationships among genes. GRAVITY differs from existing methods by integrating gene regulatory relationships, thereby enabling the simultaneous, systems-level prediction of kinetic parameters for all genes. Furthermore, GRAVITY enables the characterization of cellular heterogeneity in transcriptional dynamics by predicting each cell kinetic parameters without any additional constraints on *α*_*i j*_, *β*_*i j*_, and *γ*_*i j*_. Thus, the kinetic parameters can be formulated as follows:

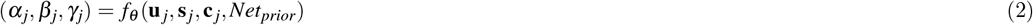

Where *α*_*j*_, *β*_*j*_ and *γ*_*j*_ denotes the vectors of all genes’ kinetic parameters in cell *j*. The vectors **u**_*j*_ and **s**_*j*_ denote the unspliced and spliced abundances of all genes in the cell *j*, respectively. The vector **c**_*j*_ denotes the cellular embeddings. In GRAVITY, we directly use the UMAP embeddings as the cellular embeddings to preserve the cellular heterogeneity and ensure consistency between the inferred velocities and the visualization. *Net*_*prior*_ is a binary directed adjacency matrix that indicates the predefined background regulatory network.

### GRAVITY model architecture

The architecture of GRAVITY is composed of three key modules: the transcriptional encoding module converts the original scalar unspliced and spliced abundances of genes into vector embeddings for preparing to decipher complex gene regulatory relationships; From there, the regulatory network inference module employs the masked multi–head cross attention to identify cell-specific regulatory relationships; Finally, the updated gene embeddings are passed to the kinetic parameter inference module to reveal the underlying transcriptional dynamics.

#### Transcriptional encoding module

Given the unspliced and spliced abundances *u*_*i j*_ and *s*_*i j*_ of gene *i* in cell *j*, previous studies, such as CellDancer, directly treat the expression values as continuous scalars. This conventional representation limits the ability of the subsequent modules to decipher gene regulatory relationships. Therefore, the transcriptional encoding module converted the expression scalars into learnable embeddings. Specifically, GRAVITY multiplies the unspliced (*u*_*i j*_) and spliced (*s*_*i j*_) values by the gene specific learnable weights 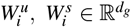 Moreover, most existing RNA velocity inference methods rely on a predefined UMAP/t-SNE plot. GRAVITY also incorporates cell coordinate information as initial cellular embeddings to enhance the cellular heterogeneity of expression representation. Finally, the scalars *u*_*i j*_ and *s*_*i j*_ were converted to the embeddings 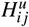 and 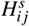 by the concatenation of expression representation and cellular representation:

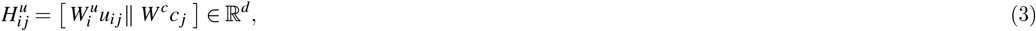

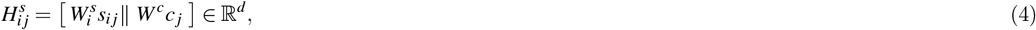

Here, *c*_*j*_ denotes the coordinate information of cell *j* or other user-provided cell representations, and *W*^*c*^ is a learnable transformation matrix that unifies the dimensions of the initial cell representations to *d*_*c*_. ∥ denotes the concatenation operation, and *d* = *d*_*g*_ + *d*_*c*_. Hereafter, we use the simplified notation of 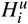 and 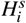 to denote the unspliced and spliced embeddings for gene *i* implicitly omitting the cellular index.

#### Regulatory network aware module

The main assumption of GRAVITY is that the transcriptional dynamics is also associated with the underlying gene regulation. Specifically, the unspliced abundance of a target gene is also affected by its upstream regulatory factors. To model the regulatory relationships, GRAVITY employs a cross-attention mechanism, where attention is defined as follows:

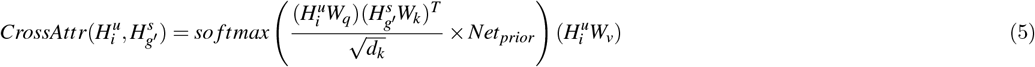

Here, *W*_*q*_, *W*_*k*_, and *W*_*v*_ are the learnable weight matrices for the query, key, and value in the attention mechanism, respectively. *g*^*′*^ dentoes the gene set without the target gene *i* and *d*_*k*_ is the dimension of the keys. To enhance the biological interpretability of the model and reduce computational complexity, the predefined background network *Net*_*prior*_ is used as a binary mask to prioritize the potential direct regulatory relationships. For instance, in the pancreatic dataset, the background network reduced the number of potential regulatory relationships from 3.5 million to 100,000, representing a 97% reduction. Our method also provides flexibility for users to specify and input their custom background network, or even without it.

#### Kinetic parameter inference module

Based on updated unspliced and spliced embedding 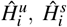 of gene *i*, GRAVITY then concatenates them and feeds them into an multi-layer perceptron (MLP) 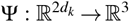 with a softplus activation function to predict kinetic parameters α_*i*_, β_*i*_ and γ_*i*_. Unlike previous inference methods, this MLP uses shared weights for all genes. To align the predicted kinetic parameters with the scale of the unspliced and spliced RNA abundance, we multiply them by three factors derived from the dataset (Supplementary Notes). Given the scaled kinetic parameters 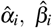 and 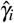 the extrapolated mRNA abundance is estimated with a small time step Δ*t* (with a default value of 0.5)

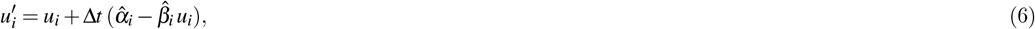

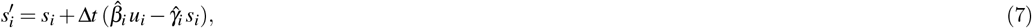

For all gene in cell *j*, we use the simplified notation of 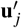 and 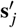 to denote the extrapolated unspliced and spliced vectors.

#### Two-stage optimization strategy and optimization objectives

GRAVITY employs a two-stage optimization strategy, with distinct optimization objectives designed for each stage: a cell-wise stage and a gene-wise stage. During the cell-wise stage, a triplet contrastive loss is employed to optimize for consistency in the overall cell velocity within a neighborhood while maximize the velocity difference with distantly located cells. Given a given cell *j*, we identify its *k*^+^ nearest neighbors as positive set and *k*^−^ farthest cells as negative set, and the triplet loss function is defined as:

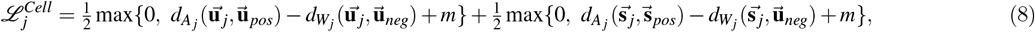

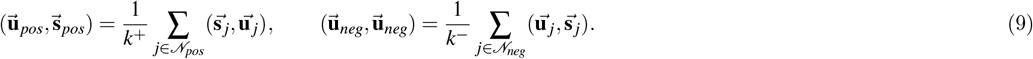

Here, the velocity vectors of cell *j* are represented by 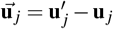 and 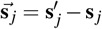. *m* is the margin and the function 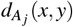 denotes the calculation of a weighted Euclidean distance. *A*_*j*_ is the vector of gene importance, derived by the weighted out-degree of each gene using cross-attention. If a gene has no out-degree, GRAVITY assigns a small predefined weight. Biologically, a gene’s impact on the inference of cell velocity is greater when it serves as a crucial regulatory factor for that cell.

After cell-wise stage training, GRAVITY establishes the coherent cell-level velocities. Then, in the gene-level stage, the global cell-level velocity serves as a foundational reference to guide the optimization of each gene’s velocity. Using the estimated cell velocity 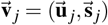 and its corresponding embedding *c*_*j*_ of cell *j*, GRAVITY projects the cell velocity into the cell embedding space to generate an initial pesudo cell embedding 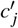, which represents the its future state. This pseudo cell is then mapped to a real cell within its local neighborhood, which is defined as follows:

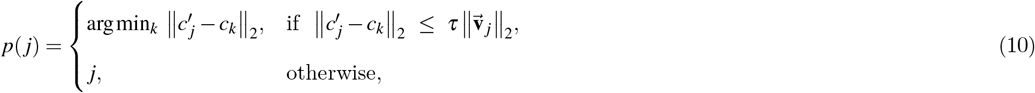

This mapping strategy enables the cell velocity to more accurately reflect the intrinsic manifold of the dataset, while also providing a reliable reference for optimizing the kinetic parameters of each gene. Furthermore, for cells located at the developmental boundary where no neighboring real cell is identified, GRAVITY directs the cell’s future state towards itself, thereby mitigating the appearance of abnormal cell velocities. Finally, the gene-wise loss function is defined as follows:

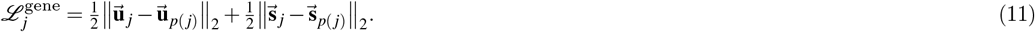

Unlike the cell-wise loss function, the gene-wise loss function equitably minimizes the discrepancy between the predicted and the real unspliced and spliced mRNA abundance for each gene. In addition, the cell-wise training stage involves training the entire model, while during the gene-wise training phase, we freeze all weights except for the last three layers of the kinetic parameter inference module. This strategy accelerates training efficiency, while the fine-tuned gene kinetic parameters further optimize the overall cellular velocity.

#### Branching-aware trajectory consistency metric

CBDir and ICCoh are two of the most representative metrics to quantify visual assessment of the plotted velocity^8, 10, 48^. CBDir assesses the developmental trajectory of cells at cross-cell states boundaries by quantifying the alignment between its inferred cell velocity and its expected path within a shared UMAP space. Meanwhile, ICCoh focuses on the directional consistency of cell velocity within a given cluster by averaging the cell coherence scores, which are calculated using a cosine similarity function in a cell’s local neighborhood. However, both metrics provide only a partial view of cell velocity performance, either from the perspective of directional accuracy or neighborhood consistency. For instance, while ICCoh reflects neighborhood consistency, it fails to capture the accuracy of velocity direction. Furthermore, the validity of CBDir inherently depends on the accurate UMAP localization of cells with distinct states. Inspired by prior studies^3, 49^ on cell trajectory analysis, we propose a Branching-aware Trajectory Consistency (BATC) metric to comprehensively evaluate inferred cell velocities.

For a predefined lineage *A* → *B*, we first fit a smooth principal curve *γ*_*A*→*B*_(*u*) on the UMAP embedding of *A* ∪ *B*, project each cell *c* ∈ *I*_*A*∪*B*_ to its closest point *p*(*c*) on the curve, and use the local tangent 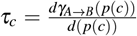 as the reference direction. The edge-wise alignment is

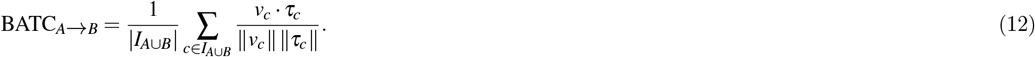

To account for branching, for each source cluster *A* with outgoing targets Out(*A*) = {*B*_1_, …, *B*_*m*_}, we compute the per-cell cosine scores on every outgoing edge,

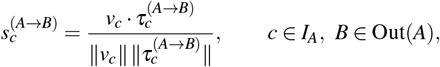

and retain, for each *c* ∈ *I*_*A*_, the best-matching branch by taking the maximum across edges:

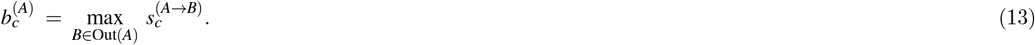

The dataset-level BATC is then reported as the cell-weighted mean of these branch-aware scores over all source clusters involved in the predefined graph:

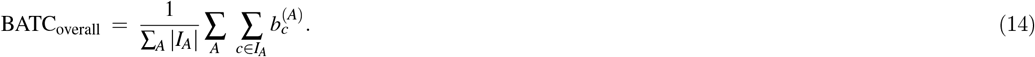

Here, *I*_*A*_ and *I*_*A* ∪ *B*_ denote the cell sets belonging to cluster *A* and *A* ∪ *B*, respectively; *v*_*c*_ is the predicted RNA velocity in the embedding; 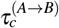 is the tangent on γ_*A*→*B*_ at the projection of *c*; and the maximization is taken across all outgoing edges of the same source *A*. In practice we guard against zero-norm vectors and use nanmean in the above averages.

#### Regulatory gene importance scoring

We utilize the masked attention matrix *M*_*attr*_ as the inferred regulatory network for each cell to assess the importance of regulatory genes. Specifically, for each target gene, we first identify its top 10% most influential regulatory genes as the key regulators based on their attention scores. These key regulators are then consolidated into a comprehensive set. The final importance score for each regulatory gene is calculated by summing its regulatory influence across all target genes it regulates.

Furthermore, with the regulatory gene importance scores for all cells, we perform the differential analysis to identify cell type-specific regulatory genes. Regulatory genes exhibiting significant differential importance are defined as cell type-specific regulatory genes and then ranked based on their *LogFC* values.

#### Pathway importance scoring

We also assess the importance of a biological pathway using the inferred regulatory network *M*_*attr*_. A higher pathway importance score indicates an increased level of pathway activity within the cell. For a given biological pathway, we identify the intersections of its regulatory genes and target genes with those in the attention matrix *M*_*attr*_, denoted as *S* and *T*, respectively. The importance within the cell is then determined by summing the weights of the regulatory relationships between regulatory genes and target genes, defined as follows:

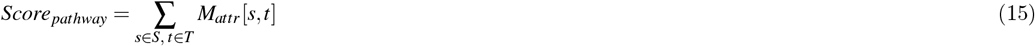

#### Cell embedding learning

Unlike the original cell embeddings, we obtained the RNA velocity-driven cellular embeddings by averaging the unspliced and spliced expression embeddings across all genes. These two embeddings were then concatenated to represent the final representation for that cell. This learned cell embedding provides a more accurate representation of cell-to-cell similarity based on RNA velocity. When visualized by UMAP, these embeddings exhibit a strong concordance with the underlying RNA velocity field.

#### In silico perturbation experiments

To dissect the regulatory mechanisms governing cell development, we proposed an in silico strategy to simulate alterations in cell velocity resulting from perturbations of regulatory genes. This perturbation experiment involves two types of stimulation: inhibition and activation. We first leverage unperturbed single-cell RNA sequencing (scRNA-seq) data to train a model and establish a baseline cell velocity landscape. For inhibition experiments, both the unspliced and spliced expressions of the inhibited gene in specific cell types are set to zero, and concurrently masking its regulatory relationships within the attention mechanism. Conversely, gene activation was modeled by significantly overexpressing the target gene in a cell-type-specific manner. The resulting perturbed expression profiles are then used to re-infer cellular velocities with the trained model. By comparing the cell velocity before and after the perturbation, we can directly assess the impact of these alterations on inferred cell trajectories and developmental fate decisions.

#### Regulatory module enrichment analysis

To systematically dissect the regulatory programs within a given cell type, we constructed regulatory modules from the top 15 most important regulatory genes and their corresponding top 3 target genes, as determined by the masked attention matrix *M*_*attr*_. Then, We performed GO enrichment analysis using ClusterProfiler^50^ and pathway enrichment analysis with Enrichr^51–53^, with the results then visualized using ActivePathways^47^, Cytoscape^54^, and EnrichmentMap^55^.

#### GRN validation

To validate the accuracy of inferred regulatory relationships by GRAVITY, we leveraged reference ChIP-seq data for specific regulatory factors from Cistrome DB^38^ and ChIP-seq Atlas^56^ as the gold standard. A regulatory relationship was considered valid if the predicted target gene exhibited a significant binding signal in the corresponding cell-type-specific ChIP-seq data.

## Code availability

The GRAVITY tool is developed as a Python package and is openly available for use at https://github.com/CSUBioGroup/GRAVITY.git.

## Data availability

All scRNA-seq and scATAC-seq datasets used in this study are publicly accessible. Preprocessed datasets have been deposited to Zenodo and can be accessed at: https://doi.org/10.5281/zenodo.17626914.

## Acknowledgements

This work has been supported by the National Natural Science Foundation of China under Grant (No.62225209, No.62572492). The results here are in part based upon data generated by the TCGA Research Network: https://www.cancer.gov/tcga and the National Cancer Institute Clinical Proteomic Tumor Analysis Consortium (CPTAC).

## Author contributions

Z.M., R.Z. and M.L. conceived and supervised the study. Z.M. designed and implemented the GRAVITY framework and conducted case studies under the supervision of R.Z. and M.L.. Z.M., Z.F. and R.Z. analysed the results and drafted the manuscript. R.Z. and M.L. provided critical revisions of the manuscript. X.S. and Y.Z. contributed to the design and execution of multi-omics downstream analyses, and T.W. assisted with language editing and grammatical revision of the manuscript. All authors reviewed and approved the final version of the manuscript.

