## Supplemental Files for "GRAVITY: Dynamic gene regulatory network-enhanced RNA velocity modeling for trajectory inference and biological discovery"

### Supplementary Figures

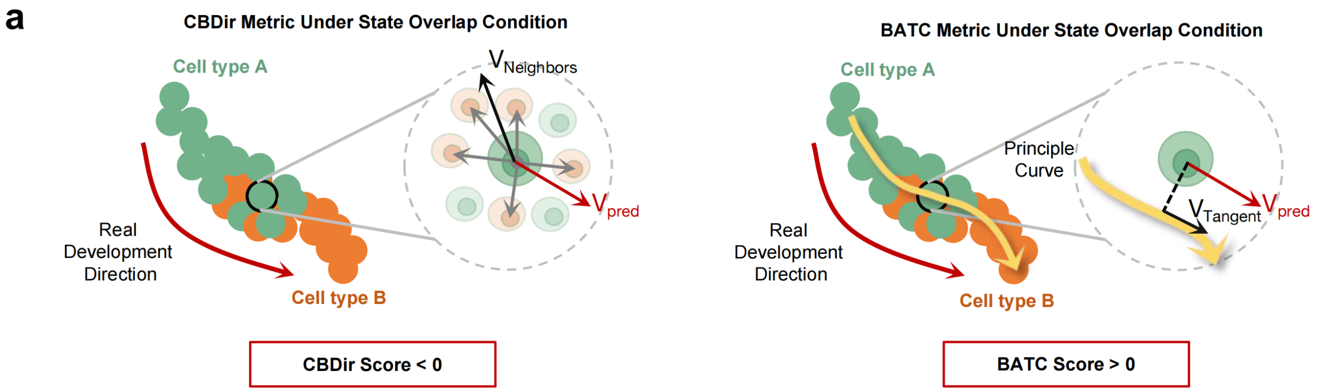


**Supplementary Figure S1. Schematic of the two-stage optimization strategy and the BATC metric. a,** Schematic of the CBDir and BATC metrics within inter-cell-type boundary regions. Upper panel: In the mixed boundary regions, CBDir may erroneously assign negative scores to the correct developmental direction, owing to the mislocalization of next-state cells. Lower panel: BATC utilizes the principal curve of the predefined cell lineage as the reference direction, effectively mitigating the bias introduced by cell mislocalization.

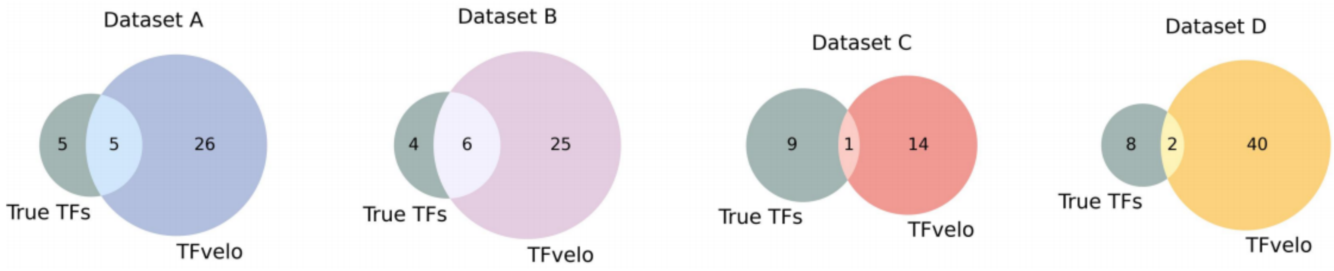


**Supplementary Figure S2. TF overlap between true TFs and genes selected by TFvelo on different datasets.** Venn diagram illustrating the overlap between TFs from the prior network and the rate genes selected by TFvelo in the simulated dataset.

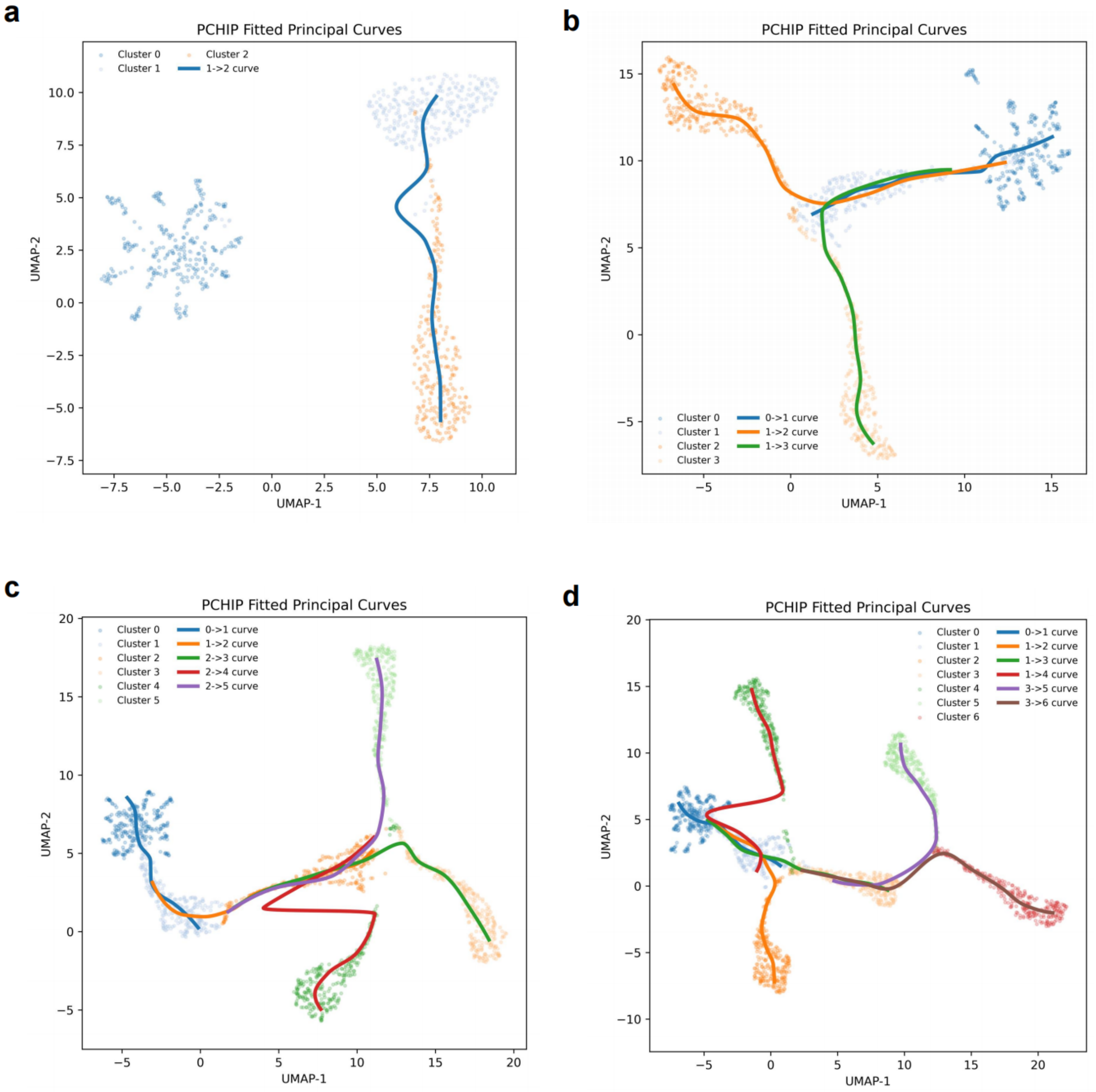


**Supplementary Figure S3. Fitted principal curves plot of the BATC metric in the simulated dataset. a-d**, Principal curves fitted across the four simulated datasets, with different colors indicating distinct ground-truth transition directions.

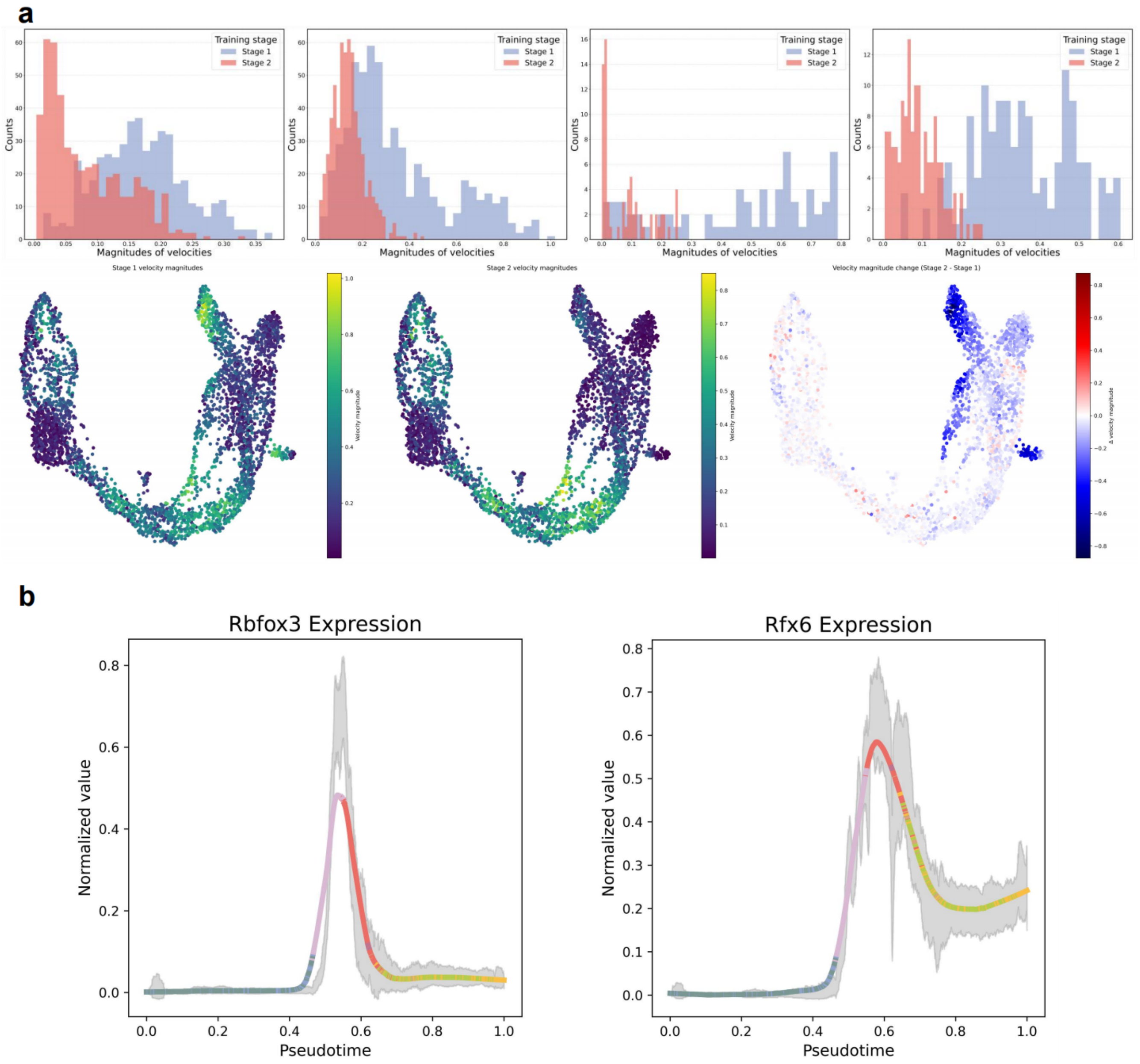


**Supplementary Figure S4. Comparative analysis of cell velocities inferred by GRAVITY two-stage optimization in pancreatic development. a**, Comparison of cell velocity magnitudes obtained by the cell-wise and gene-wise stages in four mature cell types. Upper panel: distributions of velocity magnitudes for Alpha, Beta, Delta and Epsilon cell. Lower panel: heatmap of velocity magnitudes during pancreatic development, showing (left to right) cell velocity magnitudes after cell-wise training, magnitudes after gene-wise training, and the difference between the two metrics, respectively. **b**, Gene expression dynamics of $Rbfox3$ and $Rfx6$ along the inferred pseudotime, with different colors indicating cell types.

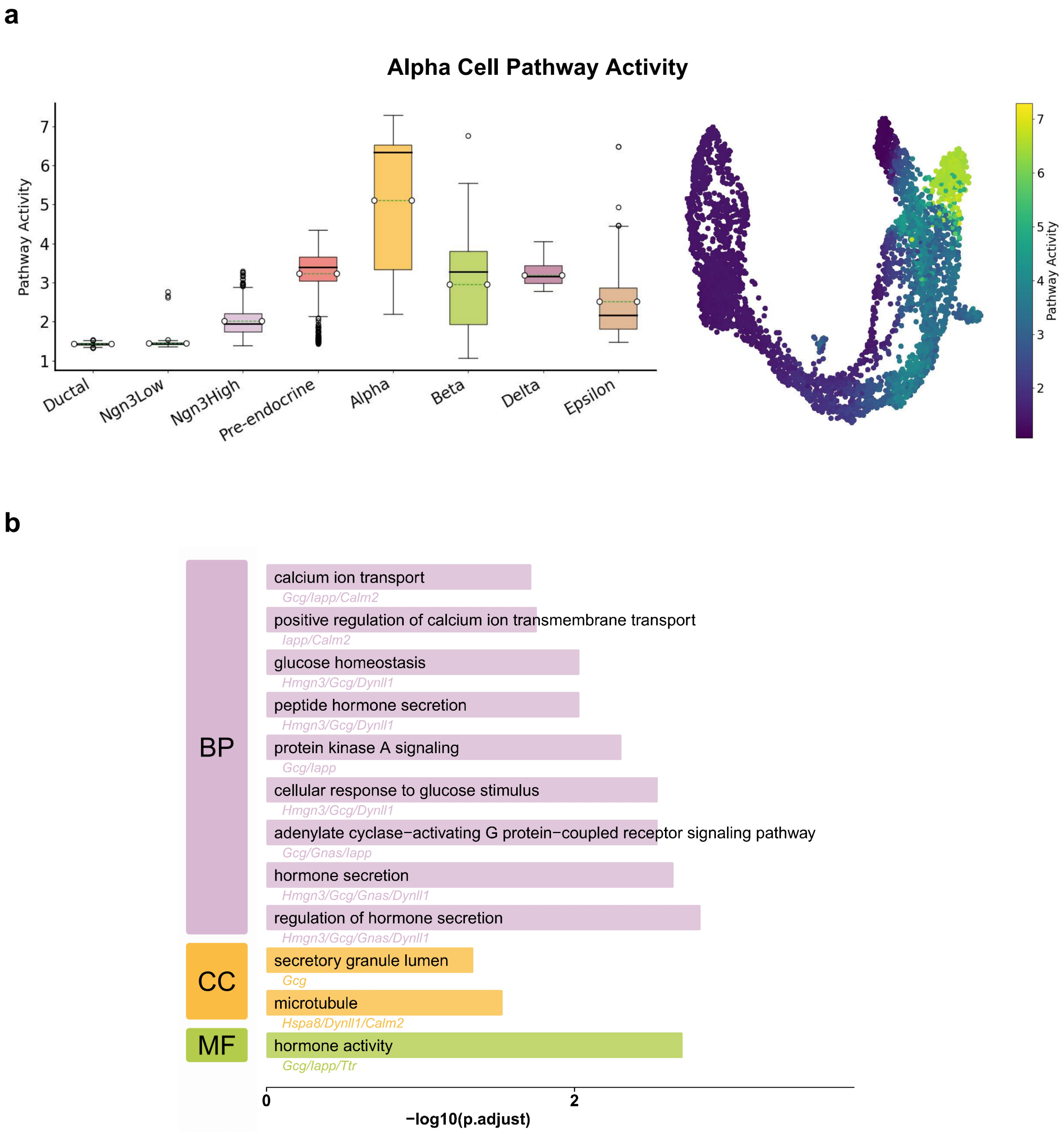


**Supplementary Figure S5. Alpha cell pathway activity and GO enrichment analysis based on pathway genes and top regulatory genes identified by GRAVITY. a**, Visualization of the alpha cells pathway activity scores across different cell types. **b**, GO enrichment analysis of top 15 genes identified by GRAVITY in alpha cells.

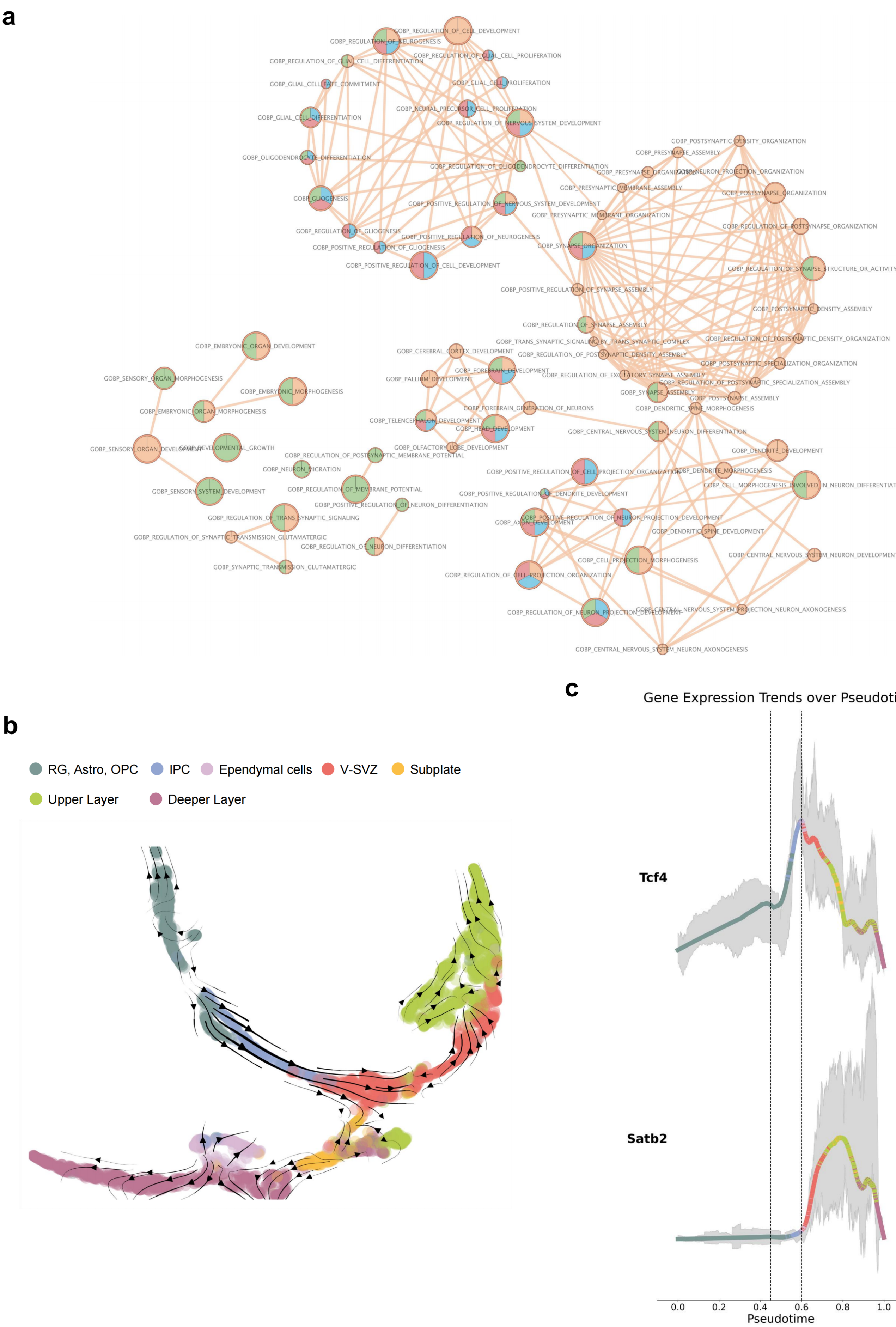
a
GOBP_REGULATION_OF_CELL_DEVELOPMENT
GOBP_REGULATION_OF_NEUROGENESIS
GOBP_REGULATION_OF_GLIAL_CELL_PROLIFERATION
GOBP_REGULATION_OF_GLIAL_CELL_DIFFERENTIATION
GOBP_GLIAL_CELL_FATE_COMMITMENT
GOBP_GLIAL_CELL_PROLIFERATION
GOBP_NEURAL_PRECURSOR_CELL_PROLIFERATION
GOBP_GLIAL_CELL_DIFFERENTIATION
GOBP_REGULATION_OF_NERVOUS_SYSTEM_DEVELOPMENT
GOBP_OLIGODENDROCYTE_DIFFERENTIATION
GOBP_REGULATION_OF_OLIGODENDROCYTE_DIFFERENTIATION
GOBP_GLIOGENESIS
GOBP_POSITIVE_REGULATION_OF_NERVOUS_SYSTEM_DEVELOPMENT
GOBP_REGULATION_OF_GLIOGENESIS
GOBP_POSITIVE_REGULATION_OF_NEUROGENESIS
GOBP_POSITIVE_REGULATION_OF_GLIOGENESIS
GOBP_POSITIVE_REGULATION_OF_CELL_DEVELOPMENT
GOBP_POSTSYNAPTIC_DENSITY_ORGANIZATION
GOBP_PRESYNAPSE_ASSEMBLY
GOBP_PRESYNAPSE_ORGANIZATION
GOBP_NEURON_PROJECTION_ORGANIZATION
GOBP_PRESYNAPTIC_MEMBRANE_ASSEMBLY
GOBP_POSTSYNAPSE_ORGANIZATION
GOBP_PRESYNAPTIC_MEMBRANE_ORGANIZATION
GOBP_REGULATION_OF_POSTSYNAPSE_ORGANIZATION
GOBP_SYNAPSE_ORGANIZATION
GOBP_REGULATION_OF_SYNAPSE_STRUCTURE_OR_ACTIVITY
GOBP_POSITIVE_REGULATION_OF_SYNAPSE_ASSEMBLY
GOBP_POSTSYNAPTIC_DENSITY_ASSEMBLY
GOBP_REGULATION_OF_SYNAPSE_ASSEMBLY
GOBP_TRANS_SYNAPTIC_SIGNALING_BY_TRANS_SYNAPTIC_COMPLEX
GOBP_REGULATION_OF_POSTSYNAPTIC_DENSITY_ORGANIZATION
GOBP_REGULATION_OF_POSTSYNAPTIC_DENSITY_ASSEMBLY
GOBP_POSTSYNAPTIC_SPECIALIZATION_ORGANIZATION
GOBP_REGULATION_OF_EXCITATORY_SYNAPSE_ASSEMBLY
GOBP_REGULATION_OF_POSTSYNAPTIC_SPECIALIZATION_ASSEMBLY
GOBP_SYNAPSE_ASSEMBLY
GOBP_POSTSYNAPSE_ASSEMBLY
GOBP_DENDRITIC_SPINE_MORPHOGENESIS
GOBP_EMBRYONIC_ORGAN_DEVELOPMENT
GOBP_SENSORY_ORGAN_MORPHOGENESIS
GOBP_EMBRYONIC_MORPHOGENESIS
GOBP_EMBRYONIC_ORGAN_MORPHOGENESIS
GOBP_SENSORY_ORGAN_DEVELOPMENT
GOBP_DEVELOPMENTAL_GROWTH
GOBP_SENSORY_SYSTEM_DEVELOPMENT
GOBP_CEREBRAL_CORTEX_DEVELOPMENT
GOBP_FOREBRAIN_DEVELOPMENT
GOBP_PALLIUM_DEVELOPMENT
GOBP_FOREBRAIN_GENERATION_OF_NEURONS
GOBP_TELENCEPHALON_DEVELOPMENT
GOBP_HEAD_DEVELOPMENT
GOBP_OLFACTORY_LOBE_DEVELOPMENT
GOBP_REGULATION_OF_POSTSYNAPTIC_MEMBRANE_POTENTIAL
GOBP_NEURON_MIGRATION
GOBP_REGULATION_OF_MEMBRANE_POTENTIAL
GOBP_POSITIVE_REGULATION_OF_NEURON_DIFFERENTIATION
GOBP_REGULATION_OF_TRANS_SYNAPTIC_SIGNALING
GOBP_REGULATION_OF_SYNAPTIC_TRANSMISSION_GLUTAMATERGIC
GOBP_REGULATION_OF_NEURON_DIFFERENTIATION
GOBP_SYNAPTIC_TRANSMISSION_GLUTAMATERGIC
GOBP_CENTRAL_NERVOUS_SYSTEM_NEURON_DIFFERENTIATION
GOBP_DENDRITE_DEVELOPMENT
GOBP_POSITIVE_REGULATION_OF_CELL_PROJECTION_ORGANIZATION
GOBP_DENDRITE_MORPHOGENESIS
GOBP_CELL_MORPHOGENESIS_INVOLVED_IN_NEURON_DIFFERENTIATIO
GOBP_POSITIVE_REGULATION_OF_DENDRITE_DEVELOPMENT
GOBP_POSITIVE_REGULATION_OF_NEURON_PROJECTION_DEVELOPMENT
GOBP_AXON_DEVELOPMENT
GOBP_DENDRITIC_SPINE_DEVELOPMENT
GOBP_CENTRAL_NERVOUS_SYSTEM_NEURON_DEVELOPMENT
GOBP_CELL_PROJECTION_MORPHOGENESIS
GOBP_REGULATION_OF_CELL_PROJECTION_ORGANIZATION
GOBP_REGULATION_OF_NEURON_PROJECTION_DEVELOPMENT
GOBP_CENTRAL_NERVOUS_SYSTEM_PROJECTION_NEURON_AXONOGENESIS
GOBP_CENTRAL_NERVOUS_SYSTEM_NEURON_AXONOGENESIS
b
RG, Astro, OPC
IPC
Ependymal cells
V-SVZ
Subplate
Upper Layer
Deeper Layer
c
Gene Expression Trends over Pseudotime
Tcf4
Satb2
0.0
0.2
0.4
0.6
0.8
1.0
Pseudotime

**Supplementary Figure S6. Biological variation captured by GRAVITY during embryonic brain development.** **a**, Multivariate GO enrichment analysis of identified regulatory modules across four cell types (Radial Glia, IPC, Deeper Layer, and Upper Layer), where each node represents a pathway and colors indicate different cell types. **b** Visualization based on GRAVITY embeddings shows a clear branching between the Upper and Deeper Layers, and indicates putative L5 and L6 terminal states within the Deeper Layer. **c** Gene-expression dynamics of *Tcf4* and *Satb2* along GRAVITY inferred pseudotime. Smoothed lines trace the mean trend, grey bands show variability across cells, and vertical dashed lines mark the inflection points of gene expression for both genes along pseudotime.

| Gene | LogFC | p-value |
|---|---|---|
| INS2 | 143.99306 | 6.923081290304737e-50 |
| HMGN3 | 43.44871 | 1.4611772125352027e-22 |
| IAPP | 20.58693 | 4.106267282392726e-185 |
| FOS | 15.94245 | 7.815558179863369e-23 |
| PDX1 | 13.94042 | 2.2933303162594704e-250 |
| NNAT | 12.70526 | 7.62967616105133e-89 |
| GNAS | 11.97134 | 7.296437203276326e-57 |
| TTR | 6.74027 | 1.9963534738752404e-78 |
| DYNLL1 | 6.35203 | 7.669523649738851e-80 |
| TUBA1A | 6.17288 | 1.0313339492583824e-176 |
| CALM2 | 5.57794 | 5.664063715254681e-45 |
| CALR | 5.55448 | 7.48212229055806e-80 |
| REST | 5.47458 | 3.3907541993477435e-29 |
| PYY | 5.02762 | 1.8189282834340666e-145 |
| HSPA5 | 4.63327 | 3.6731404104928655e-189 |
| SEC61B | 3.52618 | 1.2174306002654962e-254 |
| RBP4 | 3.06436 | 1.1741994563483413e-128 |
| EZH2 | 3.05874 | 9.481059409051314e-07 |
| IGSF3 | 2.92317 | 5.209954057473259e-28 |
| ARRDC4 | 2.80026 | 0.8009541350274392 |
| CHGB | 2.75501 | 6.032355936500466e-102 |
| TRIM47 | 2.70355 | 3.0061983430102757e-243 |
| TTYH1 | 2.62125 | 2.487014474075173e-59 |
| DLK1 | 2.58531 | 1.0553094026245163e-236 |
| MOSPD1 | 2.56046 | 6.514404078878678e-165 |
| CCND1 | 2.27645 | 2.222348072203032e-62 |
| RAP1B | 2.12972 | 1.1634403113726826e-149 |
| GNG12 | 1.66400 | 2.198991900906372e-146 |
| PDIA6 | 1.62467 | 1.152939786906743e-164 |
| TESC | 1.50849 | 0.0005468124552708212 |

**Supplementary Table S1. Important genes identified by GRAVITY in Beta cells and their LogFC scores and significance.**

| Gene | Literature |
|---|---|
| Pyy | PYY-Dependent Restoration of Impaired Insulin and Glucagon Secretion in Type 2 Diabetes following Roux-En-Y Gastric Bypass Surgery |
| Nkx2-2 | Nkx2.2-repressor activity is sufficient to specify -cells and a small number of -cells in the pancreatic islet |
| St18 | Myt Transcription Factors Prevent Stress-Response Gene Overactivation to Enable Postnatal Pancreatic b Cell Proliferation, Function, and Survival |
| Pclo | Alteration in glucose homeostasis and persistence of the pancreatic clock in aged mPer2$^{Luc}$*mice* |
| Sphkap | Sphingosine kinase 1-interacting protein is a novel regulator of glucose-stimulated insulin secretion |

**Supplementary Table S2. Genes and references supporting an indirect association with Pdx1 or relevance to pancreatic islet beta cells within the identified Pdx1 module.**

| Retain Gene | | | | |
|---|---|---|---|---|
| TP53 | OLR1 | CD83 | ICAM1 | CXCR4 |
| CXCR1 | CXCR2 | PTGS2 | SELL | CSF3R |
| FCGR3B | AGO4 | ARG1 | CYP4F3 | ERGIC1 |
| FLOT2 | FRAT2 | LRP10 | MGAM | MMP25 |
| MSRB1 | NDEL1 | NFE2 | PADI4 | PBX2 |
| PHOSPHO1 | RASGRP4 | REPS2 | SULT1B1 | TSEN134 |
| XKR8 | CCR3 | CCRL2 | DDIT3 | FLOT1 |
| HIF1A | IRAK2 | MAFF | MAP1LC3B2 | MCCLN1 |
| NBN | NOD2 | PI3 | PLAU | PPIF |
| TGM3 | TOM1 | UBR5-AS1 | ZNF267 | WNT4 |
| WNT9A | WNT2B | WNT3A | WNT11 | WNT10B |
| WNT5B | WNT3 | WNT6 | WNT10A | WNT7B |
| WNT7A | WNT5A | WNT5A-AS1 | WNT2 | WNT16 |

**Supplementary Table S3. List of retain genes for HVG selection of NSCLC neutrophil dataset.**

### Supplementary Notes

**Classical estimation via analytic/approximate solutions.**
We take $t \geq 0$ to denote an *elapsed time interval* from a local anchor $t_0$, i.e., we write $u_{ig}(t) := u_{ig}(t_0 + t)$ and $s_{ig}(t) := s_{ig}(t_0 + t)$ with initial conditions $u_{ig}(0) = u_{ig}(t_0)$ and $s_{ig}(0) = s_{ig}(t_0)$. Over any interval on which the rates are piecewise constant, the standard first–order kinetics

$$\frac{du_{ig}}{dt} = \alpha_{ig} - \beta_{ig} u_{ig}, \qquad \frac{ds_{ig}}{dt} = \beta_{ig} u_{ig} - \gamma_{ig} s_{ig}, \tag{1}$$

admit closed–form *interval solutions* $(u_{ig}(t), s_{ig}(t))$. For constant $(\alpha_{ig}, \beta_{ig}, \gamma_{ig})$ on an interval of length $t$,

$$u_{ig}(t) = \Big(u_{ig}(0) - \tfrac{\alpha_{ig}}{\beta_{ig}}\Big) e^{-\beta_{ig} t} + \tfrac{\alpha_{ig}}{\beta_{ig}}, \tag{2}$$

$$s_{ig}(t) = e^{-\gamma_{ig} t} s_{ig}(0) + \frac{\beta_{ig}\Big(u_{ig}(0) - \tfrac{\alpha_{ig}}{\beta_{ig}}\Big)}{\gamma_{ig} - \beta_{ig}} \Big(e^{-\beta_{ig} t} - e^{-\gamma_{ig} t}\Big) + \tfrac{\alpha_{ig}}{\gamma_{ig}}\big(1 - e^{-\gamma_{ig} t}\big). \tag{3}$$

Classical RNA–velocity pipelines do not solve for $(\alpha, \beta, \gamma)$ in closed form; instead, they *estimate* these rates by fitting snapshot data to relations derived from (2)–(3) (e.g., the near–steady regression $\beta u \approx \gamma s$ to infer $\gamma/\beta$ after fixing a global scale), or by fitting on/off closed forms with latent switching times via likelihood/EM. These strategies rely on parametric trajectory shapes and inherit time–scale non–identifiability (joint rescaling of the interval $t$ and $(\alpha, \beta, \gamma)$ leaves the mean trajectories invariant), typically handled by fixing one degree of freedom or normalizing time.

**Velocity embedding**
We project the high-dimensional per-cell velocity vectors onto the fixed 2D manifold. Let $C = [c_1, \dots, c_N]^\top \in \mathbb{R}^{N \times d}$ be the embedding with $c_j \in \mathbb{R}^d$ (typically $d = 2$). For cell $j$, let $\mathbf{s}_j \in \mathbb{R}^G$ denote the spliced-expression vector and $\mathbf{s}'_j \in \mathbb{R}^G$ the *extrapolated* spliced abundances obtained from the kinetic update (Kinetic parameter inference module, step size $\Delta t$). The high-dimensional velocity is

$$\vec{\mathbf{s}}_j \;=\; \mathbf{s}'_j - \mathbf{s}_j \;\in\; \mathbb{R}^G. \tag{4}$$

***Neighbors.*** Build a $k$-nearest-neighbors graph $A \in \{0,1\}^{N \times N}$ (either in the embedding $\{c_j\}$ or in expression space $\{\mathbf{s}_j\}$), with $A_{jj} = 0$ and degree $\deg(j) = \sum_k A_{jk}$.

***Forward-alignment scores.*** Compare $\vec{\mathbf{s}}_j$ with the expression displacement to each neighbor $k$, $\mathbf{d}_{jk} = \mathbf{s}_k - \mathbf{s}_j$. We compute the *Pearson correlation* across genes:

$$r_{jk} \;=\; \frac{\big(\mathbf{d}_{jk} - \bar{\mathbf{d}}_{jk}\big)^\top \big(\vec{\mathbf{s}}_j - \bar{\vec{\mathbf{s}}}_j\big)}{\big\|\mathbf{d}_{jk} - \bar{\mathbf{d}}_{jk}\big\|_2 \, \big\|\vec{\mathbf{s}}_j - \bar{\vec{\mathbf{s}}}_j\big\|_2}, \qquad r_{jj} = 0. \tag{5}$$

***Transition weights.*** Map the similarity scores to a row-stochastic transition over the $k$-NN using an exponential kernel with *kernel width* $\sigma > 0$ (default $\sigma = 0.05$):

$$P_{jk} \;=\; \frac{\exp\big(r_{jk}/\sigma\big) A_{jk}}{\sum_\ell \exp\big(r_{j\ell}/\sigma\big) A_{j\ell}}, \qquad P_{jk} \geq 0, \; \sum_k P_{jk} = 1. \tag{6}$$

Here, $\sigma$ controls the kernel's selectivity: smaller $\sigma$ yields a *narrow* kernel that concentrates mass on the most aligned neighbors (sharper weighting), while larger $\sigma$ yields a *broader* kernel that distributes mass more evenly (softer averaging).

***Unit directions on the embedding.*** Define the unit displacement from $j$ to $k$ in the embedding:

$$u_{jk} = \begin{cases} \dfrac{c_k - c_j}{\|c_k - c_j\|_2}, & k \neq j, \\ \mathbf{0}, & k = j. \end{cases} \tag{7}$$

***Projected velocity.*** Aggregate neighbor directions with correlation-induced weights $P_{jk}$, then subtract the *uniform* ($A_{jk}$-only) mean over the $k$-NN to remove geometric border drift:

$$\vec{\mathbf{v}}_j^{(\text{emb})} = \sum_k P_{jk} u_{jk} - \frac{1}{\deg(j)} \sum_k A_{jk} u_{jk} = \sum_k \left( P_{jk} - \tfrac{A_{jk}}{\deg(j)} \right) u_{jk}.$$

#### Dataset-derived scaling of kinetic parameters

Here we details how the three dataset-derived scaling factors are computed and applied to the MLP outputs $\alpha_i, \beta_i, \gamma_i$ to obtain the scaled parameters $\hat{\alpha}_i, \hat{\beta}_i, \hat{\gamma}_i$ used in the kinetic update.

***Per-gene reference statistics.*** For each gene $i$, let $u_{ji}$ and $s_{ji}$ denote the unspliced and spliced abundances in cell $j$ (after the same normalization used by the model). Compute per-gene maxima over cells:

$$u_{\max,i} = \max_j u_{ji}, \qquad s_{\max,i} = \max_j s_{ji}. \tag{8}$$

***Scaling factors.*** Guided by the steady-state identities $u^* = \alpha/\beta$ and $s^* = \alpha/\gamma$, define three nonnegative, per-gene factors

$$\alpha_i^0 = 2u_{\max,i}, \qquad \beta_i^0 = 1, \qquad \gamma_i^0 = \frac{u_{\max,i}}{s_{\max,i}}. \tag{9}$$

***Applying the scales.*** Let $\alpha_i, \beta_i, \gamma_i \geq 0$ be the softplus-transformed outputs of the shared MLP $\Psi$ (main text). The scaled parameters are obtained by per-gene broadcasted multiplication:

$$\hat{\alpha}_i = \alpha_i \cdot \alpha_i^0, \qquad \hat{\beta}_i = \beta_i \cdot \beta_i^0, \qquad \hat{\gamma}_i = \gamma_i \cdot \gamma_i^0. \tag{10}$$

*(i)* $\alpha_i^0 = 2u_{\max,i}$. With $\beta_i^0 = 1$ fixing the time unit, $u^* \approx \alpha/\beta$ suggests $\alpha$ should be on the order of $u$. We choose $2 \times u_{\max,i}$ to (a) match the observed unspliced range and (b) provide headroom for ongoing transcriptional bursts, avoiding systematic underestimation when many cells are near steady-state. *(ii)* $\beta_i^0 = 1$. This pins the global time scale to the update step $\Delta t$ and removes one free scale, simplifying optimization while leaving $\beta_i$ to capture gene- and cell-specific variation. *(iii)* $\gamma_i^0 = u_{\max,i}/s_{\max,i}$. Since $s^* \approx \alpha/\gamma$, we set $\gamma \approx \alpha/s^*$. Approximating $s^*$ by the observed $s_{\max,i}$ yields $u_{\max,i}/s_{\max,i}$ (a conservative choice that avoids overly large $\gamma$ and stabilizes the spliced update $\beta u - \gamma s$).

#### Branching-aware trajectory consistency (BATC)

Given a directed transition $A \to B$, we evaluate whether per-cell RNA velocities align with the *local trajectory direction* on the $A \cup B$ backbone in the embedding. Let $C = [c_1, \ldots, c_N]^\top \in \mathbb{R}^{N \times d}$ be the embedding ($d = 2$ by default), and let $\vec{\mathbf{v}}_j^{(\text{emb})} \in \mathbb{R}^d$ denote the velocity vector of cell $j$ in the same space. Denote the index sets $I_A = \{j : c_j \text{ in } A\}$, $I_B = \{j : c_j \text{ in } B\}$, and $I_{A \cup B} = I_A \cup I_B$. Cluster centroids are

$$\mu_A = \frac{1}{|I_A|} \sum_{j \in I_A} c_j, \qquad \mu_B = \frac{1}{|I_B|} \sum_{j \in I_B} c_j, \qquad \hat{d} = \frac{\mu_B - \mu_A}{\|\mu_B - \mu_A\|_2}. \tag{11}$$

***Edge-wise principal curve on $A \cup B$.*** We form a smooth backbone $\gamma_{A \to B} : [0,1] \to \mathbb{R}^d$ using only cells in $I_{A \cup B}$: (i) obtain a coarse A→B ordering by the 1D projection $s_j = (c_j - \mu_A)^\top \hat{d}$ and sort $I_{A \cup B}$ by $s_j$; (ii) split the range of

$\{s_j\}$ into $K$ bins and compute *skeleton points* as the mean of $c_j$ within each non-empty bin; (iii) parameterize the skeleton by normalized arc-length $u \in [0,1]$; (iv) fit a shape-preserving spline (PCHIP) to $(u,x)$ and $(u,y)$ to obtain $\gamma_{A\to B}(u) = (f_x(u), f_y(u))$. We orient the curve from $A$ to $B$ by flipping $u \mapsto 1-u$ if $(\gamma(1)-\gamma(0))^\top(\mu_B - \mu_A) < 0$.

***Closest-point projection and tangent.*** Each cell $j \in I_{A\cup B}$ is projected to its nearest point on the curve:

$$\hat{u}_j \;=\; \arg\min_{u\in[0,1]} \|c_j - \gamma_{A\to B}(u)\|_2, \qquad \tau_j \;=\; \gamma'_{A\to B}(\hat{u}_j) \in \mathbb{R}^d, \tag{12}$$

where $\tau_j$ is the *local tangent* (from the spline derivative or a small centered finite difference). The tangent orientation follows A→B by construction.

***Per-cell cosine alignment.*** We compare $\vec{\mathbf{v}}_j^{(\mathrm{emb})}$ with $\tau_j$ using a cosine score:

$$s_j^{(A\to B)} \;=\; \frac{\vec{\mathbf{v}}_j^{(\mathrm{emb})} \cdot \tau_j}{\left\|\vec{\mathbf{v}}_j^{(\mathrm{emb})}\right\|_2 \|\tau_j\|_2} \in [-1,1], \quad j \in I_{A\cup B}, \tag{13}$$

with zero-norm guarding (treated as NaN in aggregation).

***Edge-level aggregation.*** The edge-level BATC for $A \to B$ averages per-cell scores over $A \cup B$:

$$\mathrm{BATC}_{A\to B} \;=\; \frac{1}{|I_{A\cup B}|} \sum_{j\in I_{A\cup B}} s_j^{(A\to B)}. \tag{14}$$

***Branch-aware per-cell aggregation (max over outgoing edges).*** To account for branching, for each source cluster $A$ with outgoing targets $\mathrm{Out}(A) = \{B_1, \ldots, B_m\}$, we retain for every $j \in I_A$ the best-matching branch:

$$b_j^{(A)} \;=\; \max_{B\in\mathrm{Out}(A)} s_j^{(A\to B)}, \qquad j \in I_A. \tag{15}$$

The dataset-level BATC is then the cell-weighted mean of these branch-aware scores across all sources:

$$\mathrm{BATC}_{\mathrm{overall}} \;=\; \frac{1}{\sum_A |I_A|} \sum_A \sum_{j\in I_A} b_j^{(A)}. \tag{16}$$

Here, $s_j^{(A\to B)}$ is always computed with respect to the edge-specific curve $\gamma_{A\to B}$ and its local tangent at $j$'s closest-point projection; the maximization in (15) is restricted to edges sharing the same source $A$.
